# Cryptic starter amidation in antibiotic biosynthesis by *trans*-acyltransferase polyketide synthases

**DOI:** 10.64898/2026.04.24.720606

**Authors:** Yu Zhang, Marianne Costa, James A. Duncan, Lona M. Alkhalaf, Gregory L. Challis

**Affiliations:** Department of Chemistry, University of Warwick, Coventry CV4 7AL, UK; Department of Biochemistry and Molecular Biology, Biomedicine Discovery Institute, Monash University, Clayton, Victoria 3800, Australia; ARC Centre of Excellence for Innovations in Peptide and Protein Science, Monash University, Clayton, Victoria 3800, Australia

## Abstract

Polyketide biosynthesis is typically initiated by loading a starter unit onto an acyl carrier protein (ACP). In type I modular polyketide synthases (PKSs), responsible for the assembly of diverse bioactive metabolites in bacteria, this ACP is usually incorporated into a chain initiation module, alongside a starter unit loading domain. In several cases, the starter unit undergoes structural modification prior to the initiation of chain assembly. Gladiolin, an antibiotic with promising activity against bacterial and fungal pathogens, is assembled by a *trans*-acyltransferase (AT) PKS in *Burkholderia gladioli*. It appears to incorporate a succinyl starter unit, but the gladiolin PKS lacks a conventional loading module, making it unclear how this happens. The gladiolin biosynthetic gene cluster encodes an AT of unassigned function (GbnB), an ACP (GbnA), and an asparagine synthetase homolog (GbnC) with similarity to enzymes that amidate the malonyl-ACP starter unit in glutarimide antibiotic biosynthesis. Here, we elucidate a cryptic starter unit amidation mechanism in gladiolin biosynthesis involving these three proteins. GbnB loads a succinyl unit onto the phosphopantetheinyl arm of GbnA, which is subsequently amidated by GbnC using glutamine as the nitrogen donor. After the fully assembled polyketide chain is released, GbnM hydrolyzes the amide, yielding mature gladiolin. Phylogenetic analyses, coupled with gene cluster reannotation revealed analogous enzymatic machinery likely responsible for cryptic succinamyl and malonamyl starter unit incorporation into etnangien and sorangicin A, respectively. Retro-biosynthetic analyses suggest succinamyl and malonamyl starter units may be involved in the assembly of other metabolites, such as the sorangiolides and azumamides.

## Introduction

Polyketides are a large group of structurally diverse and biologically active natural products, many of which are used clinically as antibiotics, immunosuppressants and antifungals, e.g. erythromycin A, rapamycin, and amphotericin B, respectively.^1^ They are biosynthesized by polyketide synthases (PKSs), which assemble short acyl units into complex hydrocarbon scaffolds. Type I modular PKSs are giant multienzymes that typically employ a chain initiation module and a series of chain elongation modules to catalyze the sequential elongation of a starter unit with several (alkyl)malonyl extender units, accompanied by various structural modifications, resulting in the assembly of a structurally complex polyketide chain.^2,3^ Chain elongating modules contain a core set of acyl carrier protein (ACP), acyltransferase (AT) and ketosynthase (KS) domains, alongside optional domains that modify the growing polyketide chain in various ways. Type I modular PKSs are further subdivided into phylogenetically distinct *cis*-AT and *trans*-AT classes. In comparison to *cis*-AT PKSs, which contain an AT domain within each module, *trans*-AT PKSs utilize one (or occasionally more) separately encoded *trans-*acting ATs to supply extender units to each module.^4^ In addition, *trans*-AT PKSs have several other unusual features, including modules that are split across two subunits, modules that iterate (i.e. they catalyze multiple rounds of chain elongation), non-elongating ketosynthase domains,^5^ O-methylating sub-modules, and selective recruitment of *trans*-acting enzymes, such as enoylreductases, to specific modules.^4,6^

Several distinct mechanisms for chain initiation in type I modular PKSs have been reported. The most common mechanism in *cis*-AT PKSs involves an AT domain loading malonyl or methylmalonyl-coenzyme A (CoA) onto an adjacent ACP domain, followed by decarboxylation catalyzed by a KS domain variant (KS^Q^) (Figure 1A).^7^ Alternatively, an AT domain in the loading module can directly transfer the starter unit from CoA to the phosphopantetheinyl (Ppant) arm of a downstream ACP domain (e.g., loading of the propionyl starter unit in erythromycin biosynthesis).^8^ Interestingly, in this system the KS domain of the first chain elongation module can be loaded directly with the propionyl starter unit when the loading module is inactive (Figure 1B).^9^ Other well-known mechanisms for the incorporation of starter units involve acyl-ACP ligase (AL) domains, which catalyze the ATP-dependent activation of a carboxylic acid, via formation of an acyl adenylate, and subsequent acylation of the Ppant thiol in a downstream ACP domain, as observed for the loading of the 4,5-dihydroxycyclohex-1-enecarboxylic acid starter unit in rapamycin biosynthesis (Figure 1C).^10^ In this system, the resulting thioester is reduced and transferred onto the module 1 KS domain, highlighting that ACP-bound starter units can undergo structural modification prior to initiation of chain assembly.

**Figure 1.**
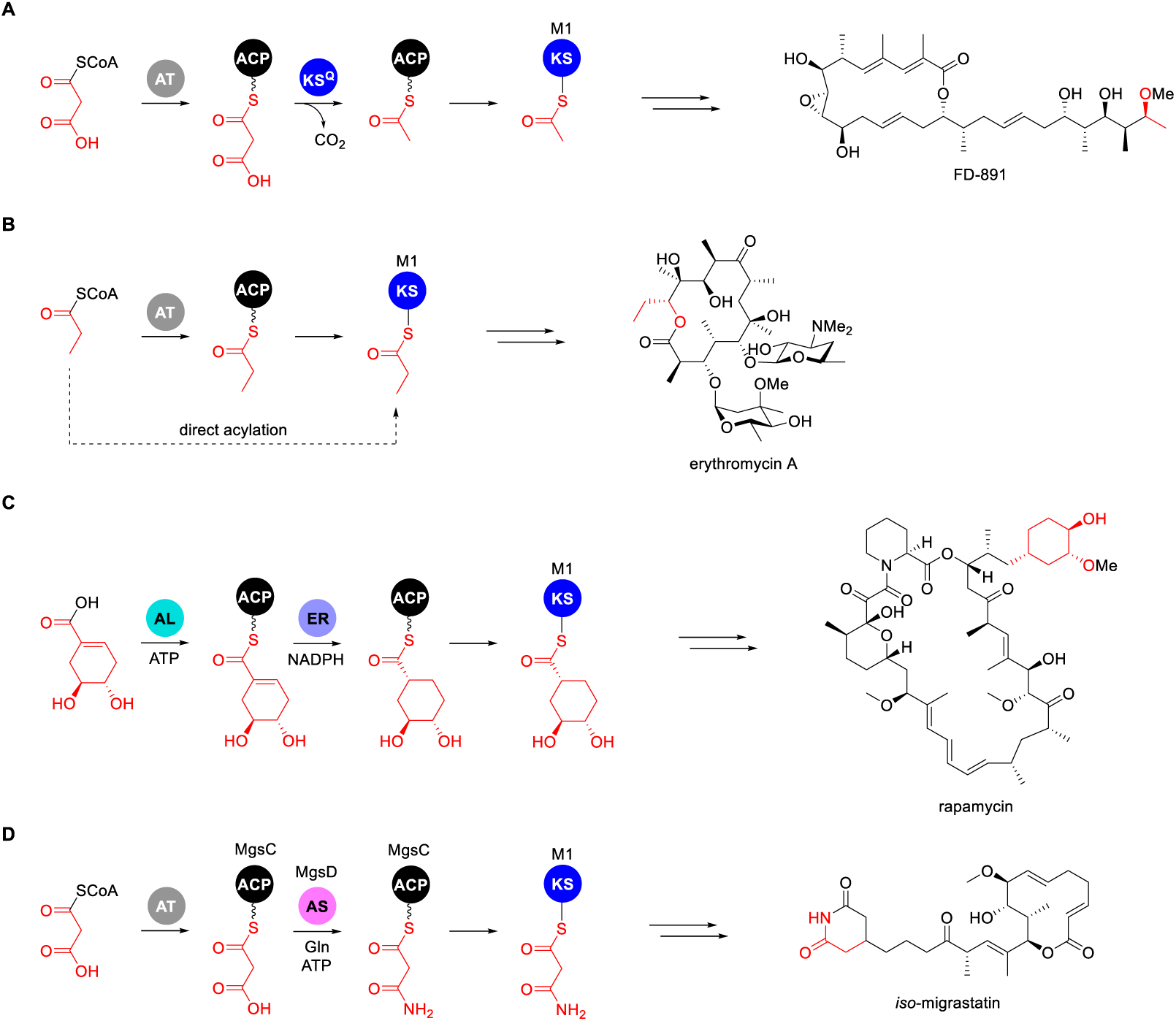
Mechanisms for chain initiation in type I modular PKSs. (A) An AT domain in the chain initiation module catalyzes malonylation of the downstream ACP domain and the upstream KS^Q^ domain catalyzes decarboxylation of the malonyl-ACP domain, thereby providing the acetyl starter unit for FD-891 biosynthesis. (B) In erythromycin biosynthesis, the propionyl starter unit is transferred by the AT domain in the chain initiation module from CoA to the downstream ACP domain. The module 1 KS domain can also accept the propionyl group directly from propionyl-CoA. (C) Rapamycin biosynthesis is initiated by ATP-dependent loading of 4,5-dihydroxycyclohex-1-enecarboxylic acid onto the ACP domain in the chain initiating module catalyzed by an upstream AL domain. The resulting enoyl thioester is reduced to the corresponding saturated thioester by an ER domain, prior to transfer of the starter unit onto the module 1 KS domain. (D) In *iso*-migrastatin biosynthesis, a malonyl group is transferred onto the chain initiating ACP by a *trans*-acting AT and is proposed to undergo amidation, catalyzed by an asparagine synthetase homolog, before being transferred to the module 1 KS domain. Starter units are highlighted in red. M1 = module 1. Domain abbreviations are as follows: AT, acyltransferase, ACP, acyl carrier protein, KS, ketosynthase (KS^Q^: KS domain variant with Cys to Gln active site mutation only able to catalyze (alkyl)malonyl-ACP decarboxylation), AL, acyl-ACP ligase, ER, enoylreductase and AS, asparagine synthetase-like amide synthetase.

Gladiolin **1** is a macrolide antibiotic produced by *Burkholderia gladioli* BCC0238, an opportunistic pathogen originally isolated from the lung of a cystic fibrosis patient (Figure 2).^11^ It has promising activity against drug resistant clinical isolates of *Mycobacterium turberculosis* and strongly potentiates the activity of amphotericin B against *Candida albicans* and other important fungal pathogens.^11,12^ Developing an in-depth understanding of gladiolin biosynthetic mechanisms will provide a foundation for biosynthetic engineering approaches to the creation of analogues, which are challenging to produce synthetically, but would provide insight into structure-activity relationships.

**Figure 2.**
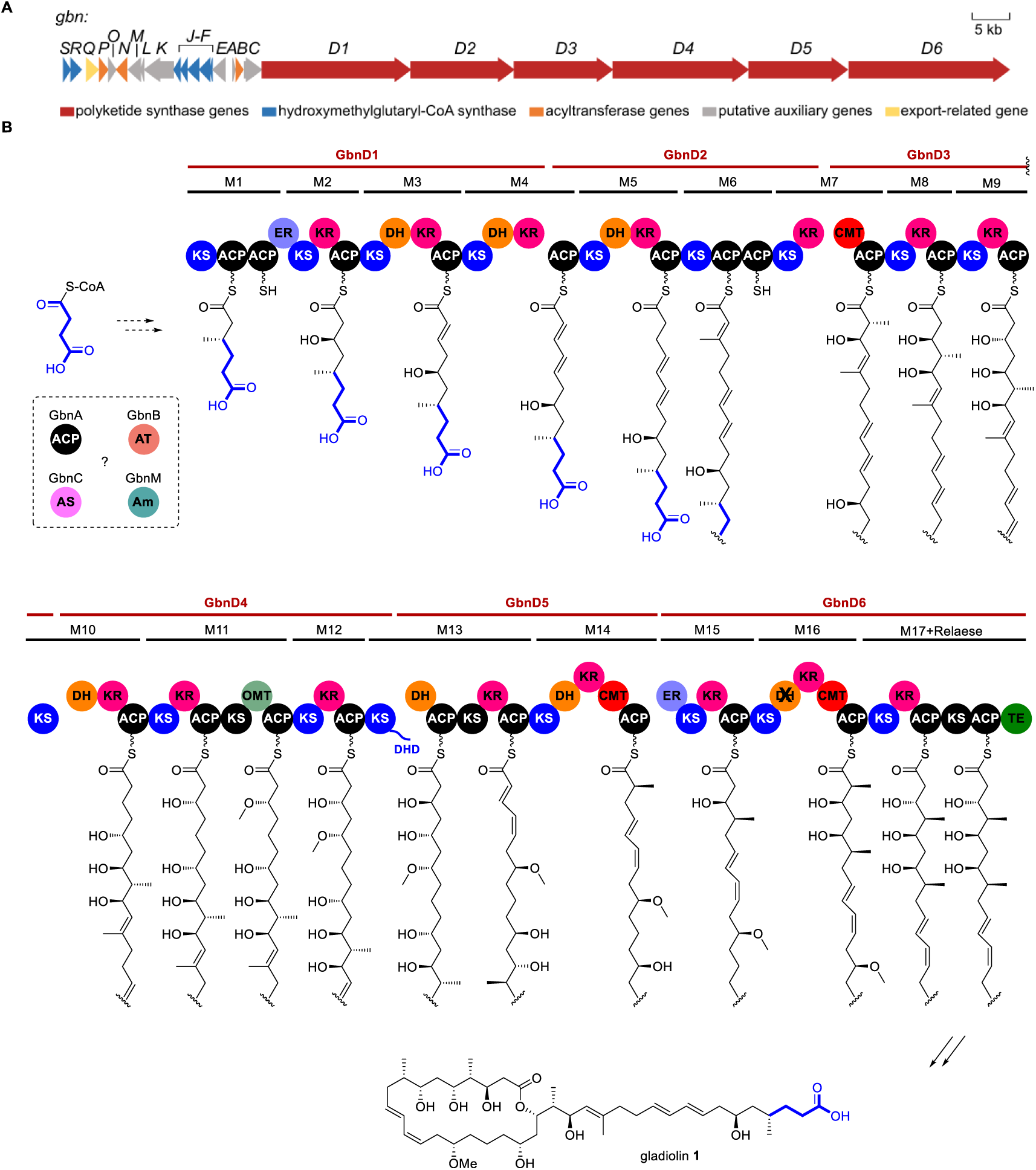
Organization of the gladiolin biosynthetic gene cluster in *B. gladioli* BCC0238 and proposed pathway for gladiolin biosynthesis. (A) The ∼106 kb *gbn* cluster consists of six large genes encoding *trans*-AT PKS subunits, and hydroxymethylglutaryl-CoA synthetase (HMGS) gene cassette for β-branch installation. (B) Previously proposed biosynthetic pathway for gladiolin. The location of the succinate-derived starter unit is highlighted in blue. The dashed box highlights putative biosynthetic enzymes encoded by the gladiolin BGC with experimentally unvalidated functions. Domain abbreviations are as follows: KS, ketosynthase (non-elongating variants are in black), ACP, acyl carrier protein, AT, acyltransferase KR, ketoreductase, DH, dehydratase, ER, enoylreductase, MT, C-/O-methyltransferase, TE, thioesterase, AS, amide synthetase and Am, amidase.

The gladiolin biosynthetic gene cluster (BGC) encodes a *trans*-AT PKS with several interesting features (Figure 2). These include an intrinsically disordered docking domain that mediates inter-subunit interaction in a split module via direct contact with a downstream catalytic domain,^13^ a dehydratase-like domain that catalyzes conversion of a (3*R*, 5*S*)-3, 5-dihydroxy thioester to the corresponding *E*, *Z*-dienoyl thioester,^14^ and a *trans*-acting enoylreductase that controls both chain length and oxidation state during chain assembly.^6^

The first subunit of the gladiolin PKS (GbnD1) lacks a canonical chain initiation module at its N-terminus for incorporation of the putative succinyl starter unit (Figure 2). This previously led us to suggest that succinyl-CoA directly acylates the module 1 KS domain,^11^ analogously to direct acylation of the corresponding KS domain in the erythromycin PKS by propionyl-CoA (Figure 1B).^9^ On the other hand, three genes of unassigned function, *gbnABC*, lie directly upstream of the PKS-encoding genes in the gladiolin BGC. These could encode proteins that play role in starter unit incorporation. Homologs of *gbnA* and *gbnC* are present in all known BGCs for the glutarimide family of polyketide antibiotics.^4,15–17^ These encode an ACP proposed to undergo malonylation (as exemplified by MgsC in *iso*-migrastatin biosynthesis) and an asparagine synthetase homolog (exemplified by MgsD) hypothesized to function as an amide synthetase (AS) that catalyzes ATP and glutamine-dependent amidation of the ACP-bound malonyl unit (Figure 1D).^18^ GbnA and GbnC are 43 and 55 % identical to MgsC and MgsD, respectively.

In addition to a canonical AT (GbnN) for loading of malonyl extender units onto ACP domains in PKS chain elongation modules and an acyl hydrolase (GbnP) for hydrolytic removal of aberrant acetyl (and potentially other short-chain acyl) groups from the PKS ACP domains,^19^ the gladiolin BGC encodes another AT-like enzyme (GbnB). A similar “additional” AT (EtnB), with 53% sequence identity to GbnB, is encoded by the BGC for etnangien, an antibiotic produced by myxobacteria that is structurally related to gladiolin and is also proposed to be assembled from a succinyl starter unit.^20^

Another gene in the gladiolin BGC (*gbnM*), further upstream of the PKS genes, encodes a protein with sequence similarity to malonamidase E2,^21^ which typically catalyzes the hydrolysis of amides to produce the corresponding carboxylic acid and ammonia. In addition to the GbnB homolog EtnB, the etnangien BGC encodes homologs of GbnM (Orf5), GbnA (EtnA) and GbnC (EtnC),^20^ suggesting these proteins collectively play a key role in installing a shared structural feature of gladiolin and etnangien.

Interestingly, despite containing homologs of the AS and ACP proteins involved in incorporation of the malonamyl starter unit into glutarimide antibiotics, neither gladiolin nor etnangien contain nitrogen. Here, we use a combination of targeted gene inactivation, isolation and structure elucidation of accumulated biosynthetic intermediates, incorporation of isotope-labelled precursors, *in vitro* assays of enzymatic function, intact protein mass spectrometry, and phylogenetic analyses to reveal that the initiation of gladiolin chain assembly involves a cryptic succinamyl starter unit that is hydrolytically cleaved to produce the corresponding carboxylic acid in the final biosynthetic step. Comparative bioinformatics analyses strongly suggest that etnangien biosynthesis proceeds via an analogous mechanism. Moreover, they indicate that sorangicin, an antibiotic produced by myxobacteria that is structurally unrelated to gladiolin and etnangien,^22^ is likely assembled by a similar mechanism involving a cryptic malonamyl starter unit. Finally, a search of the literature for other natural products containing a polyketide moiety putatively assembled from a succinamyl or malonamyl starter unit identified the sorangiolides^23^ and azumamides,^24^ respectively. Together these findings suggest the malonamyl unit may be more widely employed as a starter unit for complex polyketide biosynthesis than currently appreciated.

## Results and Discussion

### Genetic characterization of roles played by *gbnA*, *gbnB*, and *gbnC* in gladiolin assembly

To investigate whether *gbnA*, *gbnB* and *gbnC* are required for gladiolin biosynthesis, in-frame deletion mutants were constructed in *B. gladioli* BCC1622, a genetically more tractable gladiolin producer,^25^ using established methods. PCR was used to confirm each mutant had the intended genotype (Figure S1). UHPLC-ESI-Q-ToF-MS analysis of ethyl acetate extracts from BSM agar cultures of the *gbnA* and *gbnB* mutants showed gladiolin production was abolished. In *trans* complementation of the *gbnA* and *gbnB* mutants respectively with the pMLBAD vector containing *gbnA* or *gbnB* under the control of an arabinose-inducible promoter partially restored gladiolin production in both cases (Figure 3).

**Figure 3.**
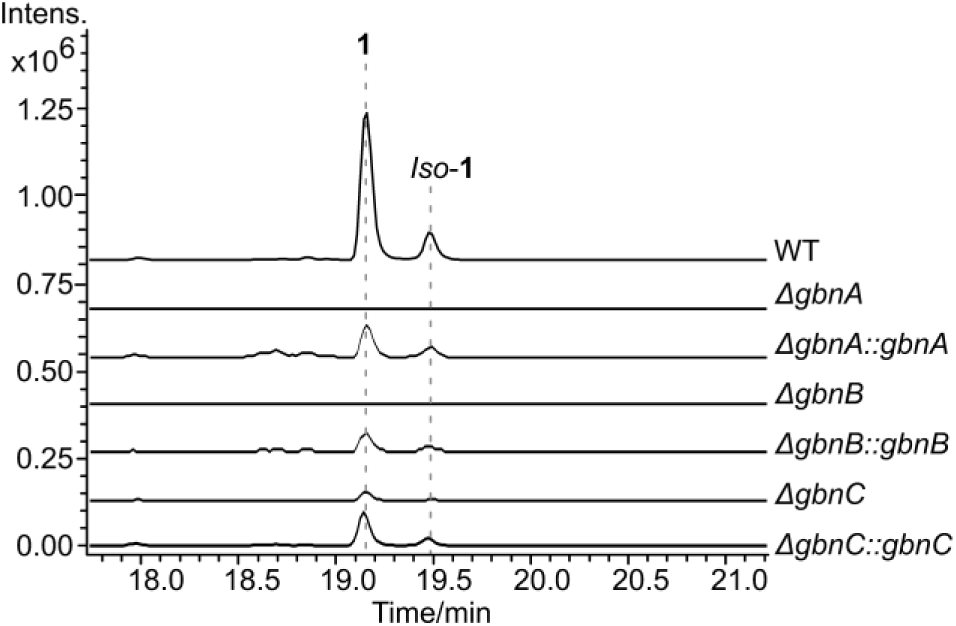
Extracted ion chromatograms at *m/z* = 801.5129, corresponding to [M+Na]^+^ for gladiolin (**1**) and its spontaneous rearrangement product *iso*-gladiolin (*iso*-**1**), from LC-MS analysis of ethyl acetate extracts of wild type (WT) *B. gladioli* BCC1622, mutants containing in-frame deletions in *gbnA*, *gbnB*, and *gbnC* and mutants complemented in *trans* with the relevant gene under the control of an arabinose-inducible promoter. The production of **1** and *iso*-**1** is abrogated in the *gbnA* and *gbnB* mutants, whereas an in-frame deletion in *gbnC* severely affects, but does not completely abolish production. In *trans* complementation of all three mutants partially restores / increases the production of **1** and *iso*-**1**.

In contrast, the production of gladiolin **1** was not completely abolished in the *gbnC* mutant, although it was severely affected (Figure 3). We hypothesize that the incomplete abrogation of production results from partial complementation by *gdsD*, a homolog of *gbnC* encoded by the BGC for gladiostatin, a polyketide antibiotic belonging to the glutarimide family also produced by *B. gladioli* BCC1622.^15^ In *trans* complementation of the *gbnC* mutant with the pMLBAD vector containing *gbnC* moderately increased gladiolin production levels.

### A novel amidated gladiolin derivative accumulates in a *gbnM* mutant

To investigate the role played by *gbnM* in gladiolin biosynthesis, in-frame deletion was introduced into this gene in *B. gladioli* BCC 1622 and PCR was used to confirm it had the intended genotype (Figure S1). UHPLC-ESI-Q-ToF-MS analysis of an ethyl acetate extract from a BSM agar culture showed gladiolin **1** production was abolished in the mutant, and a novel metabolite giving rise to ions with *m/z* = 778.5456 and *m/z* = 800.5277 was produced (Figure 4). These correspond to the [M+H]^+^ and [M+Na]^+^ ions for a compound with the molecular formula C_44_H_76_NO_10_ (calculated *m/z* = 778.5464 and 800.5284, respectively), corresponding to a derivative of gladiolin with one OH group substituted by an NH_2_ group. This novel metabolite was purified from the ethyl acetate extract of scaled-up BSM agar cultures of the *gbnM* mutant using semi-preparative HPLC and its structure was elucidated using ^1^H, ^13^C, COSY, HMBC and HSQC NMR spectroscopy (Figure 4B and S2-S6, Table S4). The ^1^H spectrum of this compound was highly similar to that reported for gladiolin.^11^ Two protons at 6.65 and 7.22 ppm, not observed in the spectrum of gladiolin and with chemical shifts characteristic of a primary amide, both correlated with C38 (*δ*_C_ 174.7 ppm) in the HMBC spectrum (Figure 4B). In addition, the proton at 6.65 ppm correlated with C37 (*δ*_C_ 32.93 ppm), confirming that the novel compound is the C38 amide derivative of gladiolin (Figure 4B), which we named gladiolamide **2**.

**Figure 4.**
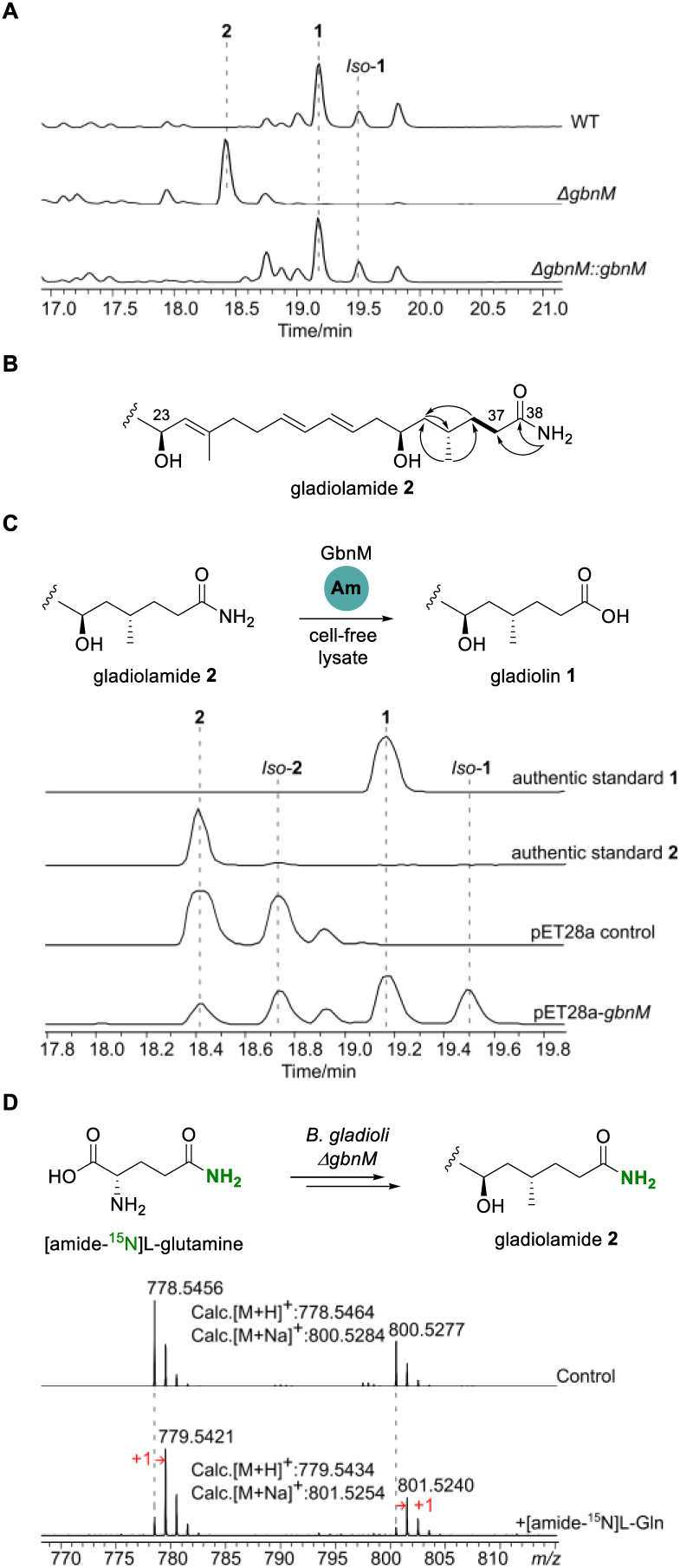
Identification and structure elucidation of biosynthetic intermediate **2** in the *gbnM* mutant of *B. gladioli* BCC1622, and confirmation of the metabolic origin of its nitrogen atom, and demonstration that GbnM has amidase activity. (A) Base peak chromatogram from UHPLC-ESI-Q-ToF-MS analyses of culture extracts from wild type *B. gladioli* BCC1622, the *gbnM* mutant, and the mutant complemented in *trans*. (B) Key HMBC and COSY correlations, represented by arrows and bold lines, respectively, that establish the location of the primary amide group in gladiolamide **2**. (C) Demonstration of GbnM amidase activity using cell-free lysates of *E. coli* BL21(DE3) carrying pET28a-*gbnM* and empty pET28a (negative control). (D) Comparison of the mass spectrum of unlabeled gladiolamide (top) and with ^15^N-labeled gladiolamide (bottom) derived from feeding of [amide-^15^N]_L_-glutamine to the *gbnM* mutant of *B. gladioli* BCC1622.

When exposed to methanol, gladiolin **1** undergoes slow ring expansion to *iso*-gladiolin *iso*-**1** the 24-membered lactone.^11^ A similar phenomenon was observed for gladiolamide **2**, resulting in the formation of an analogous 24-membered lactone isomer, which was characterized by 1- and 2-D NMR spectroscopy (Figure S7-S11, Table S5). Interestingly, the ring expansion of gladiolamide **2** to the 24-membered lactone appears to be faster than the corresponding rearrangement of gladiolin **1** (Figure S12).

### GbnM catalyzes the conversion of gladiolamide to gladiolin

The accumulation of gladiolamide **2** in the *gbnM* mutant suggested that the protein encoded by this gene catalyzes the hydrolytic conversion of **2** to gladiolin **1**. To directly test this hypothesis, we attempted to overproduce GbnM in *E. coli* BL21(DE3) as an N-terminal hexahistidine fusion protein. However, expression levels were low, making the protein challenging to purify. Thus, the catalytic activity of a cell-free lysate of *E. coli* BL21(DE3) expressing *gbnM* was investigated. UHPLC-Q-ToF-MS analysis of the cell-free lysate after addition of gladiolamide **2** showed a peak with an identical *m/z* and retention time to gladiolin **1** (Figure 4C). This was absent from a control reaction employing a cell-free lysate of *E. coli* BL21(DE3) containing the empty pET28a vector (Figure 4C). Consistent with its sequence similarity to malonamidase E2, these data indicate that GbnM catalyzes hydrolysis of the primary amide in gladiolamide **2** as the final step in gladiolin **1** biosynthesis.

### The amide nitrogen atom of gladiolamide derives from glutamine

Having established the key role played by GbnM in gladiolin **1** biosynthesis, we turned our attention to the origin of the nitrogen atom in gladiolamide **2**. The strong attenuation of gladiolin production in the *gbnC* mutant, coupled with the sequence similarity of the encoded protein to asparagine synthetase-like enzymes proposed to amidate the malonyl-ACP starter unit in glutarimide biosynthesis, led us to hypothesize that GbnC catalyzes amidation of a succinyl-ACP during gladiolin **1** starter unit assembly.

Extensive efforts to directly elucidate the function of GbnC *in vitro* were hindered by an inability to overproduce the protein in soluble form in *E. coli*. GbnC homologs encoded by glutarimide BGCs have been reported to preferentially utilize glutamine as the ammonia source for the amidation reaction.^26^ Thus, we fed [amide-^15^N]L-glutamine to BSM agar cultures of the *gbnM* mutant of *B. gladioli*. UHPLC-ESI-Q-ToF-MS analysis of ethyl acetate extracts revealed a level of ^15^N incorporation into gladiolamide **2** (Figure 4D). This confirms that the amide nitrogen atom of gladiolamide **2** is derived from glutamine and is consistent both with the proposed role of GbnC in gladiolin **1** biosynthesis, and the hypothesis that *gdsD* partially complements the *gbnC* mutant.

### Antimicrobial activity of gladiolamide

The activity of gladiolamide **2** was assessed against a range of clinically relevant bacterial pathogens, including representative members of the ESKAPE panel. In comparison to gladiolin **1**, a significant increase in activity against *Acinetobacter baumannii* and *Escherichia coli* was observed (Table 1). This enhanced activity could be attributed to an increase in permeability across the outer membrane of these Gram-negative bacteria, due to neutralization of the negatively charged C38 carboxyl group. Gladiolamide **2** showed very similar activity to gladiolin **1** against other bacteria in the ESKAPE panel and Mycobacteria. Interestingly, *iso*-gladiolamide exhibited a remarkably similar spectrum of activity to gladiolamide (Table S6).

**Table 1.**
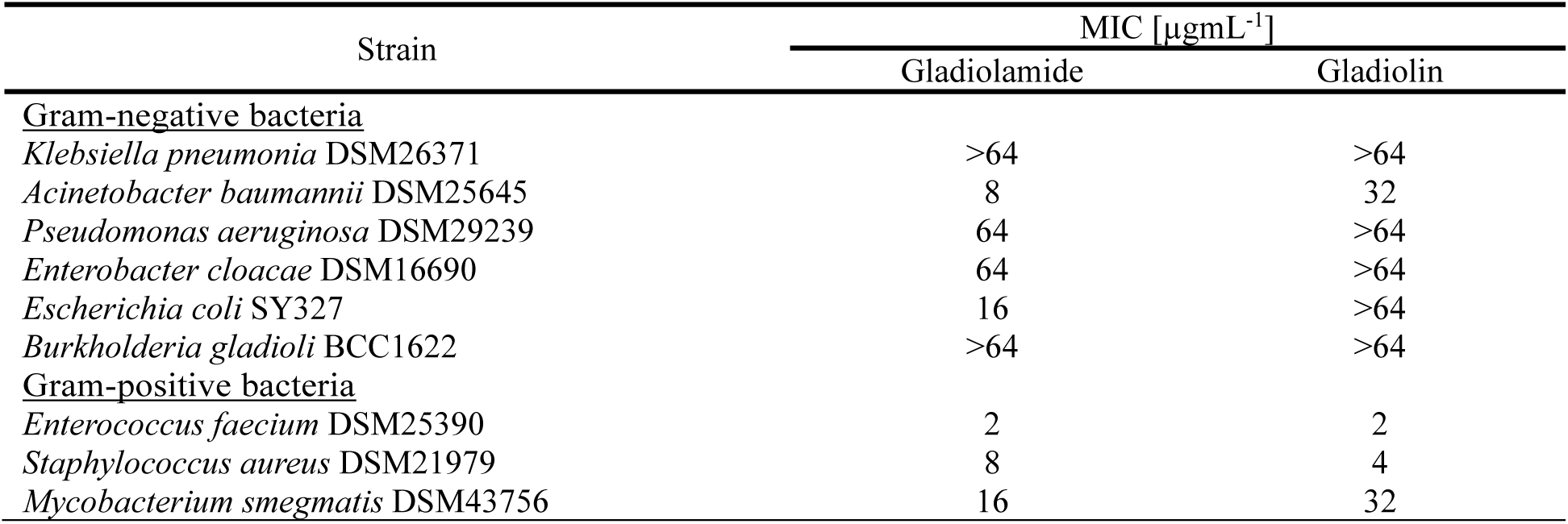
Comparison of the antibacterial activity of gladiolamide and gladiolin (MIC = minimum inhibitory concentration).

### Incorporation of ^13^C-labeled succinate into gladiolin

To establish whether the starter unit for gladiolin biosynthesis originates from succinate, we grew *B. gladioli* BCC1622 in the presence [1,4-^13^C_2_]succinic acid. UHPLC-ESI-Q-ToF-MS analysis of the culture extract showed an increase in the intensity of the ion with *m/z* = 803.5174, corresponding to [M+Na]^+^ of [^13^C_2_]gladiolin, relative to the ion with *m/z* = 801.5108, corresponding to [M+Na]^+^ of unlabeled gladiolin, compared to the relative intensity of these ions in culture extracts from *B. gladioli* BCC1622 grown in the absence of labeled succinic acid (Figure S13). Gladiolin was purified from the extract of the culture grown in the presence of labeled succinic acid using reverse phase HPLC. Comparison of the ^13^C NMR spectrum of the resulting compound with the corresponding spectrum of unlabeled gladiolin revealed a substantial increase in the intensity of the signals due to C35 and C38 in the former compared to the latter, relative to the signal due to the C15 methoxy group (Figure S13). Overall, these data demonstrate that C35-C38 of gladiolin are derived from an intact succinate unit, consistent with the hypothesis that the starter unit for gladiolin biosynthesis is derived from succinyl-CoA.

### GbnB transfers a succinyl unit onto the ACP encoded by *gbnA*

The observation that both *gbnA* and *gbnB* are essential for gladiolin biosynthesis (*vide supra*), led us to propose that the proteins encoded by these genes, along with the amide synthetase encoded by *gbnC*, together function as the chain initiation module for gladiolin biosynthesis. To investigate this hypothesis, we used intact protein mass spectrometry to investigate whether the AT encoded by *gbnB* can transfer a succinyl unit onto the ACP encoded by *gbnA*. GbnA and GbnB were overproduced as N-terminal hexa and octa-histidine fusion proteins, respectively, in *E. coli* BL21(DE3) and purified using Ni affinity chromatography (Figure S14). The identity of the purified proteins was confirmed by UHPLC-ESI-Q-ToF analysis (Figure S14). Purified GbnA was converted from the *apo*- to *holo*-form by incubation with CoA, Mg^2+^, and the phosphopantetheinyl transferase Sfp.^27^ Incubation of *holo*-GbnA with succinyl-CoA and GbnB led to the formation of succinyl-GbnA, as evidenced by a mass shift of 100 Da in the mass spectrum of the intact protein (Figure 5A). Ppant ejection from succinyl-GbnA gave rise to an ion with *m/z* = 361.1393, corresponding to succinyl-pantetheine.

**Figure 5.**
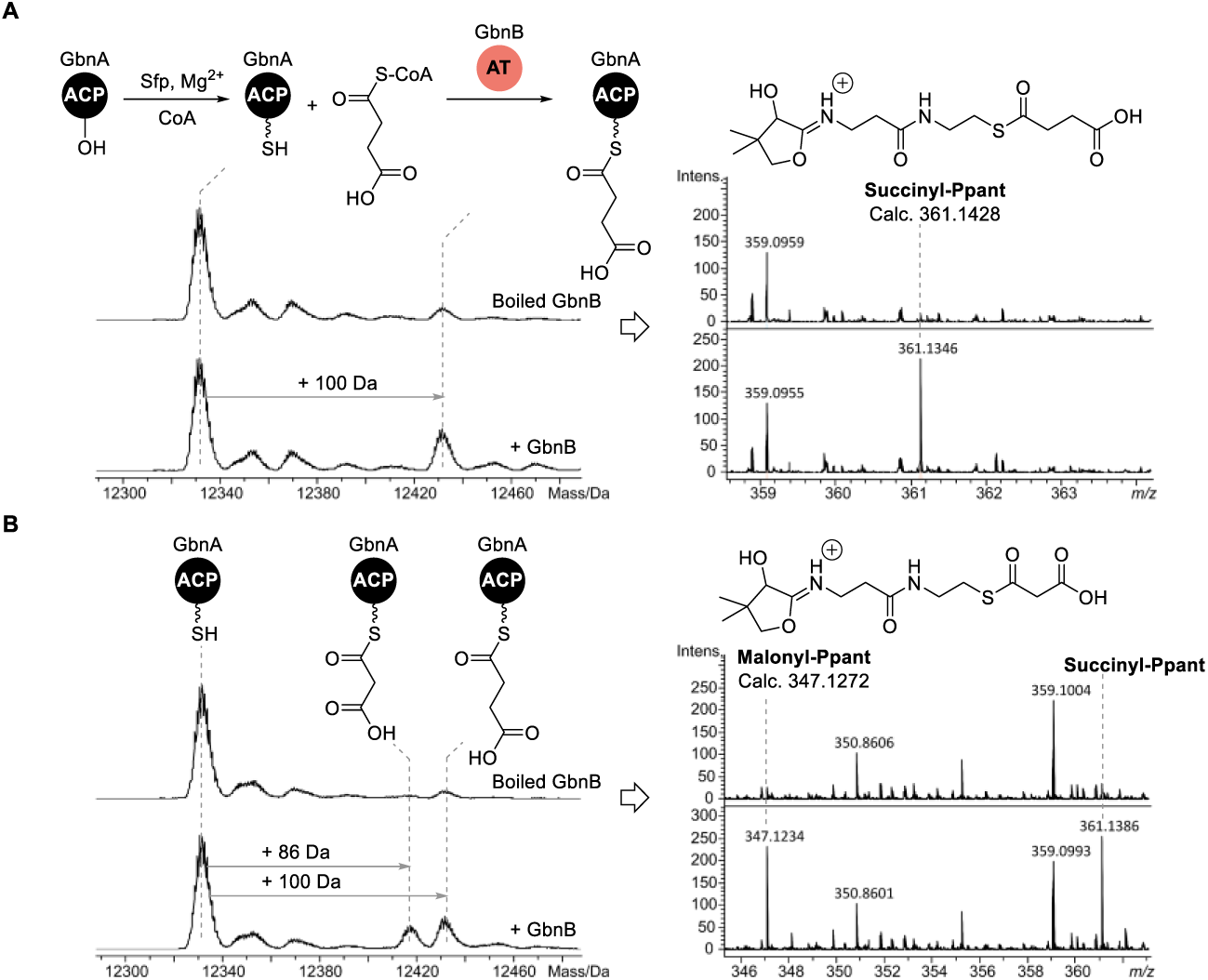
*In vitro* reconstitution of GbnB activity in transferring succinate and malonate onto *holo*-GbnA. (A) Overview of GbnB-catalyzed transacylation to *holo*-GbnA and intact protein MS analysis showing the deconvoluted mass spectra of succinyl-GbnA and Ppant ejection fragments succinyl-Ppant. (B) Deconvoluted MS spectra and Ppant ejection analysis of competition assays between malonyl- and succinyl-CoA for GbnB showing the slightly greater specificity for succinyl-CoA.

To explore the substrate specificity of GbnB, the activity of this enzyme with acetyl, malonyl and glutaryl-CoA was investigated. The formation of acetyl, malonyl, and glutaryl-GbnA was observed by intact protein MS, suggesting GbnB can tolerate succinyl-CoA analogues with varying chain lengths and terminal functionality (Figure S15). To further examine the substrate specificity of GbnB, a competition assay using equimolar amounts of succinyl and malonyl-CoA was conducted. Both intact protein MS and PPant ejection analysis of GbnA indicated a marginal preference for succinyl transfer over malonyl transfer (Figure 5B). However, succinyl thioesters are well known to undergo spontaneous cyclization to the corresponding anhydride,^28^ which may have affected the quantity of succinyl-GbnA observed in the intact protein MS analyses.

We also examined succinamyl-CoA, which could in principle be produced via direct amidation of succinyl-CoA by GbnC, as a substrate for GbnB. No formation of succinamyl-GbnA could be observed by intact protein MS (Figure S16), suggesting that GbnC-catalyzed amidation of the succinyl unit occurs after transfer onto GbnA, analogous to malonyl unit amidation in glutarimide biosynthesis.

To gain a better understanding of how GbnB differs from canonical malonyl-CoA-specific ATs that load extender units onto ACP domains in chain elongation modules in *trans*-AT PKSs, we constructed a sequence alignment. This revealed distinct differences in conserved motifs: malonyl-CoA-specific ATs typically possess GHSLG and G(A)AFH motifs, whereas the corresponding motifs in GbnB are GYSVG and GAWH (Figure S17). The significance of the H→Y, L→V, and F→W mutations in these motifs is unclear. However, the mutations are conserved in the AT encoded by *etnB* in the etnangien BGC, which is hypothesized to perform the same function as GbnB. Moreover, in Art2,^29,30^ which has been reported to catalyze succinyl transfer onto ACP domains in *trans*-AT PKSs, these motifs also contain mutations. Numerous studies that attempted to alter the substrate specificity of AT domains by mutating these conserved motifs failed to produce the expected outcomes.^31,32^ This suggests that these motifs are only partially responsible for controlling substrate specificity, as suggested by the identification of two previously unexplored motifs in the AT from the sixth chain elongation module of the erythromycin PKS via molecular dynamics simulations.^33^

### *B. gladioli* can produce a malonamide-derived gladiolamide analogue

The broad substrate tolerance of GbnB *in vitro* coupled with the observation that *gdsD* appears able to partially complement deletion of *gbnC* in *B. gladioli* BCC1622 led us to hypothesize that the gladiolin PKS may be able to assemble analogues primed with a malonamyl starter unit. Indeed, reexamination of the chromatograms from UHPLC-ESI-Q-ToF-MS analyses of the *gbnC* and *gbnB* mutant extracts identified a new species giving rise to an ion with *m/z* = 786.5115, corresponding to [M+H]^+^ for nor-gladiolamide **3** (molecular formula C_43_H_73_NNaO_10_^+^; calculated *m/z* = 786.5127) a gladiolin analogue derived from a malonamyl starter unit (Figure 6). Unfortunately, the titre of this compound was too low to permit purification and structural confirmation using NMR spectroscopy. A lower level of this metabolite was observed in extracts of the *gbnA* mutant, suggesting that GdsC, the ACP proposed to supply the malonamyl starter unit to the gladiostatin PKS,^15^ can interact productively with the first chain extension module of the gladiolin PKS (Figure 6). Very low levels of this metabolite were also observed in wild type *B. gladioli* BCC1622 (Figure 6). Interestingly, nor-gladiolin **4**, the amide hydrolysis product of nor-gladiolamide **3** was not observed, suggesting the latter is a poor substrate for GbnM.

**Figure 6.**
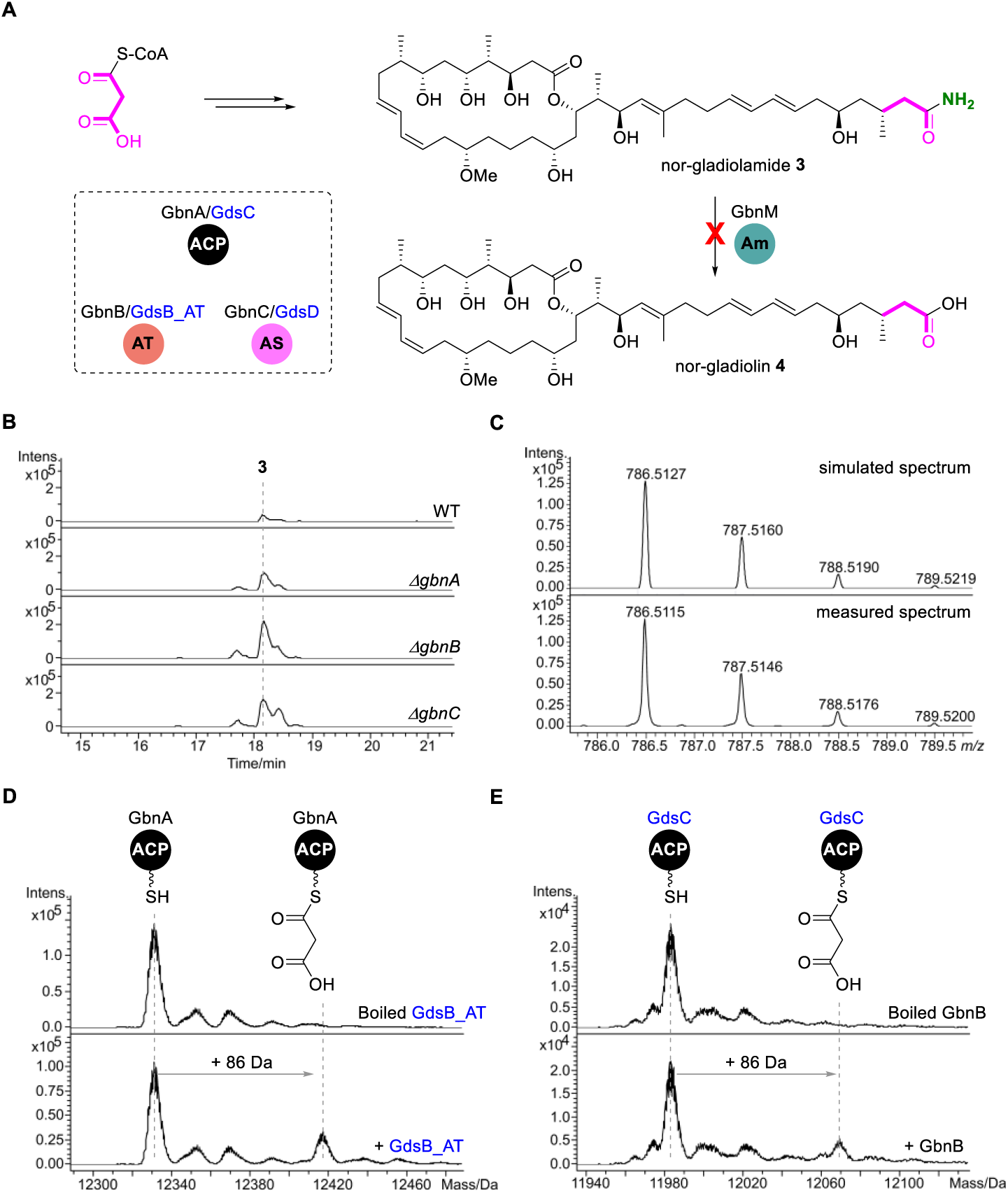
Identification of gladiolin analogues derived from a malonamyl starter unit and *in vitro* crosstalk between ACPs and ATs from the gladiolin and gladiostatin chain initiation modules. (A) Various combinations of GbnA / GdsC, GbnB / the GdsB AT domain (GdsB_AT), and GbnC / GdsD can be envisaged to provide a malonamyl starter unit to the gladiolin PKS, resulting in the production of gladiolamide / gladiolin analogues with one less carbon atom in the C21 side chain. (B) Extracted ion chromatograms at *m/z* = 786.5127 from UHPLC-ESI-Q-ToF-MS analyses of organic extracts from cultures of wild type *B. gladioli* BCC1622, and the *gbnA*, *gbnB* and *gbnC* mutants, corresponding to [M+Na]^+^ for nor-gladiolamide **3**. (C) Comparison of the simulated mass spectrum for C_43_H_73_NNaO ^+^ with the measured spectrum resulting from species eluting at 18.1 min, confirming it has a molecular formula corresponding to nor-gladiolamide **3**. (D) Crosstalk assay showing the excised GdsB AT domain catalyzes malonylation of *holo*-GbnA. (E) Crosstalk assay showing GbnB catalyzes transfer of a malonyl unit from CoA to *holo*-GdsC.

To further validate our hypothesis, we investigated whether components from the chain initiation modules of the gladiolin and gladiostatin PKSs can crosstalk *in vitro*. Intact protein MS analyses confirmed that the GdsB AT domain can malonylate GbnA and GbnB can malonylate GdsC (Figure 6). Coupled with the observation that *B. gladioli* can produce a malonamide-primed gladiolamide analogue, this highlights the potential of these components to be harnessed for biosynthetic engineering approaches to the production of novel polyketides containing malonamyl and succinamyl starter units.

Interestingly, GbnB was found to exhibit relaxed ACP specificity. In addition to malonylation of GdsC, it catalyzes succinylation of the excised ACP domain from the first chain extension module of the gladiolin PKS (Figure S18). This is consistent with a previous report that GbnB catalyzes succinylation of the excised ACP domain from the first chain extension module of the aurantinin PKS but is likely of no physiological relevance in the context of gladiolin **1** biosynthesis.^30^

### Proposed mechanism for succinyl starter unit incorporation into gladiolin

Taken together, our data suggest a plausible mechanism for net incorporation of a succinyl starter unit into gladiolin **1** (Figure 7). GbnB catalyzes transfer of a mixture of succinyl and malonyl units from CoA onto GbnA. This is reversible, as reported recently for multifunctional AT domains in fungal PKSs.^34^ The carboxyl group in the succinyl thioester undergoes selective GbnC-catalyzed amidation and the resulting succinamyl unit undergoes transacylation onto the active Cys residue of the KS domain in the first chain extension module. Assembly of the polyketide chain, followed by TE-domain catalyzed release via macrolactonization affords gladiolamide **2**. Finally, GbnM catalyzes hydrolysis of the amide to yield the mature antibiotic, gladiolin **1**.

**Figure 7.**
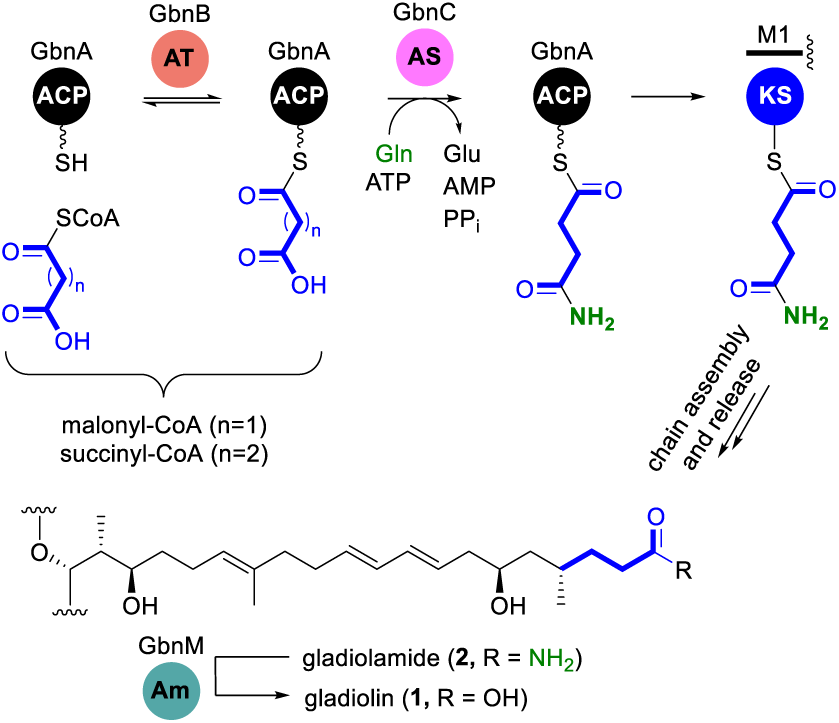
Proposed mechanism for net incorporation of a succinyl starter unit into gladiolin **1**.

### Comparative bioinformatics analyses suggest etnangien and sorangicin biosynthesis also involves cryptic starter unit amidation

Based on the presence of genes encoding homologs of GbnA, GbnB, GbnC and GbnM (*etnA*, *etnB*, *etnC*, and *orf5*, respectively) in the etnangien BGC, we hypothesize that the biosynthesis of etnangien also involves a cryptic succinamyl starter unit (Figure 8). Phylogenetic analysis of *trans*-AT PKS KS domains extracted from the MIBiG 3.0 database reveals that those from the first chain extension module in the gladiolin and etnangien PKSs clade with the corresponding KS domains in PKSs that assemble glutarimide antibiotics (Figure S19). This supports the hypothesis that succinyl-GbnA and succinyl-EtnA are amidated prior to commencement of polyketide chain assembly.

**Figure 8.**
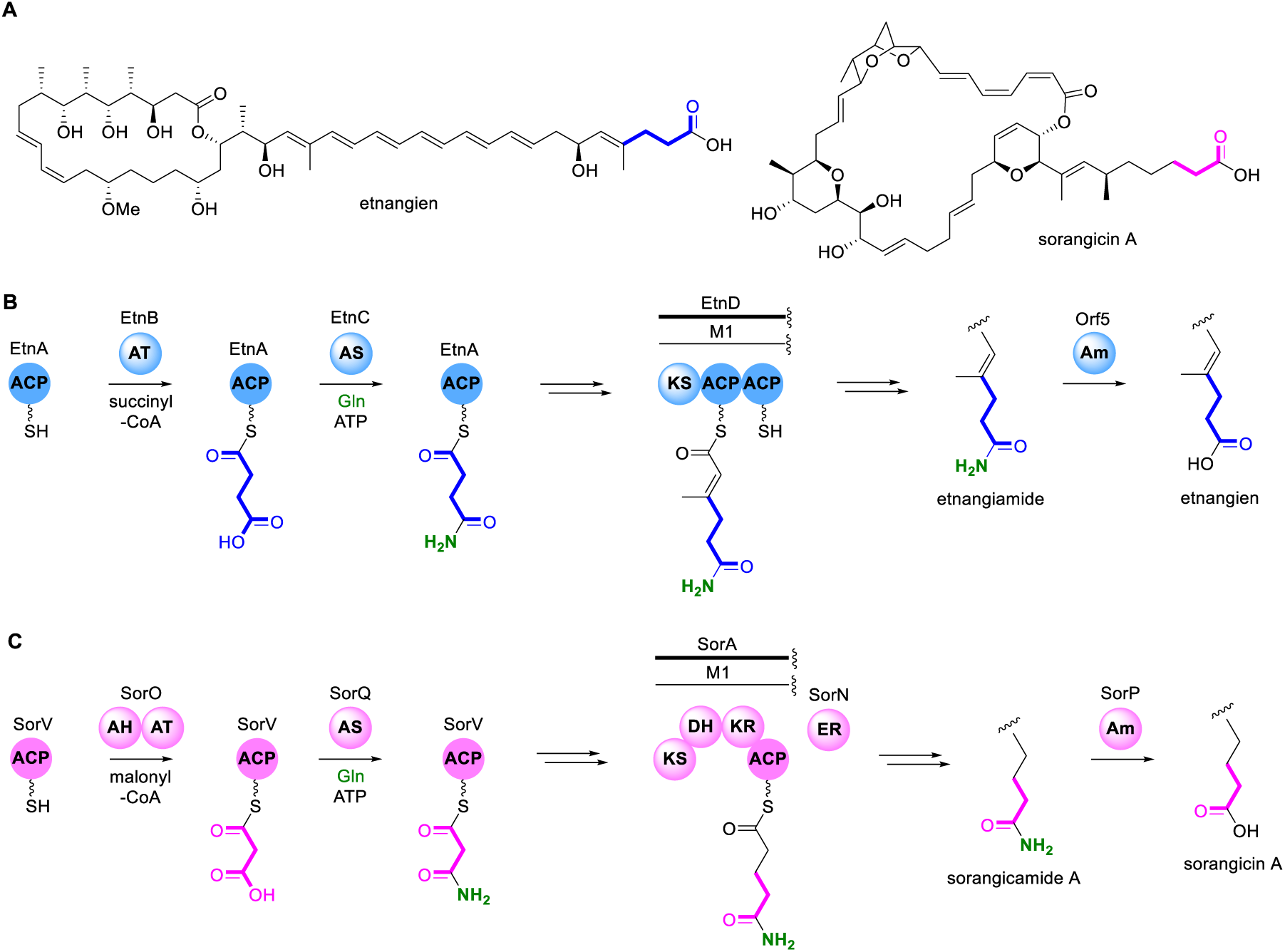
Structures of metabolites hypothesized to be assembled from cryptic sucinamyl and malonamyl starter units and proposed mechanisms for amide formation and cleavage. (A) Structures of etnangien, and sorangicin A highlighting the C and O atoms anticipated to derive from cryptic succinamyl (blue) and malonamyl (purple) starter units, respectively. Proposed mechanisms for incorporation of a succinamyl starter unit into etnangien (B) and a malonamyl starter unit into sorangicin A (C). This results in the formation of etnangiamide and sorangicamide A, putative amide derivatives of etnangien and sorangicin A, respectively, that are hydrolyzed by the amidases encoded by *orf5* and *sorP*. A dedicated AT (EtnB) loads the succinyl unit onto the ACP encoded by *etnA* in etnangien biosynthesis, whereas the C-terminal AT domain of SorO, which supplies each of the chain elongation modules with malonyl extender units, is also proposed to catalyze malonylation of the ACP encoded by *sorV* in sorangicin biosynthesis.

Sorangicin A is macrolide antibiotic produced by *Sorangium cellulosum* So ce 12 that is structurally unrelated to gladiolin and etnangien.^22^ It has previously been proposed to derive from a glutaryl starter unit.^35^ Interestingly, the KS domain from the first chain elongation module of the sorangicin PKS also clades with the KS domains from the corresponding modules of glutarimide PKSs. Furthermore, homologs of GbnC and GbnM are encoded by genes of unassigned function (*sorQ* and *sorP*) in the sorangicin BGC. This suggests that sorangicin A might be assembled from a cryptic malonamyl starter unit, rather than a glutaryl unit. However, the sorangicin BGC deposited in MiBiG lacks homologs of *gbnA / etnA / mgsC*,^35^ which encode the ACPs used for assembly of the cryptic succinamyl and malonamyl starter units in gladiolin / etnangien and glutarimide biosynthesis.

These observations prompted us to re-examine the sequence of the sorangicin BGC, which revealed a previously overlooked gene (here named *sorV*) encoding a putative ACP between *sorP* and *sorQ* (Figure S20). SorV clusters with GbnA, EtnA and homologs involved in glutarimide biosynthesis in phylogenetic sequence analyses of *trans*-AT PKS ACP domains (Figure S20). We therefore hypothesize that the biosynthesis of sorangicin A involves a cryptic malonamyl starter unit (Figure 8). The assembly of this starter unit is proposed to proceed via malonylation of SorV by the AT domain of SorO, which also supplies a malonyl extender unit to each chain elongation module in the sorangicin PKS. Amidation of malonyl-SorV is likely catalyzed by the GbnC homolog SorQ. Chain assembly and product release is proposed to result in sorangicamide, the amide analogue of sorangicin A, which is hydrolysed by the GbnM homolog SorP to yield the mature antibiotic (Figure 8).

### A potential role for amidated starter units in sorangiolide and azumamide biosynthesis

In addition to the sorangicins, *S. cellulosum* So ce 12 has been reported to produce sorangiolides A and B, macrolide antibiotics with modest activity against Gram-positive bacteria (Figure 9).^23^ Structural inspection of the sorangiolides suggest they are likely assembled by a modular PKS from a succinyl starter unit, and either twelve malonyl extender units plus six S-adenosylmethionine (SAM)-derived methyl groups, or twelve methylmalonyl-CoA extender units - depending on whether the PKS belongs to the *trans*-AT or *cis*-AT class (Figure 9). Given that *S. cellulosum* appears to employ a cryptic malonamyl starter unit for assembly of the sorangicins, it is tempting to speculate that the putative succinate-derived carbon atoms in the sorangiolides arise from a succinamyl starter unit, assembled and eventually hydrolytically cleaved via an analogous mechanism to that employed in gladiolin and etnangien biosynthesis.

**Figure 9.**
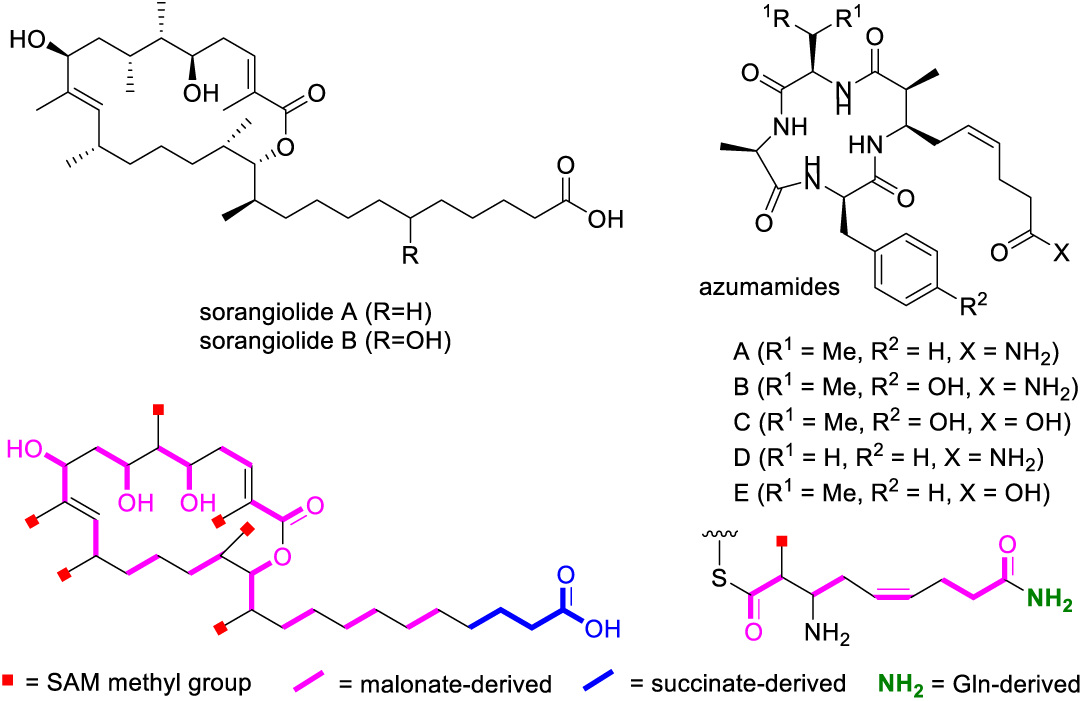
Structures of the sorangolides and the azumamides (top) and the proposed biosynthetic origins (according to *trans*-AT PKS biosynthetic logic) of the polyketide portions of the major congener in each case (bottom).

The azumamides are a family of tetrapeptides isolated from the marine sponge *Mycale izuensis* collected in Japanese coastal waters.^24^ In addition to three proteinogenic amino acids, these metabolites contain a conserved β-amino acid capped with either a carboxamide or a carboxylate group (Figure 9). The congeners containing the carboxylate capped β-amino acid have potent activity (IC_50_ = 10-70 nM) against purified recombinant HDACs 1, 2, 3, 10 and 11, whereas the corresponding carboxamides are about two orders of magnitude less active against the same enzyme panel (IC_50_ = 3-6 μM).^36^ Interestingly, when tested against total HDACs in HeLa cell extracts, the carboxamides displayed similar activity to the corresponding carboxylates, likely because the former are converted to the latter by hydrolases in the cell extracts.^37^ A synthetic analogue of azumamide C lacking the carboxylate group in the β-amino acid was two orders of magnitude less active against HDACs 1-3 than the natural product.^38^ Taken together, these observations suggest (i) that the carboxylate-capped β-amino acid is the key pharmacophore for Class I HDAC inhibition by this natural product family; (ii) the carboxylate group likely binds strongly to the catalytically essential Zn^2+^ ion in the active site of Class I HDACs; and (iii) the azumamides containing the carboxamide-capped β-amino acid, which constitute nearly 70% of all congeners isolated from the sponge, likely function as pro-drugs that are converted to their active carboxylate-capped forms by hydrolases in target cells.

Retro-biosynthetic dissection of the azumamides indicates that the carboxamide-capped β-amino acid could be assembled by a modular PKS from a malonamyl starter unit, and three malonyl extender units plus a SAM-derived methyl group, or two malonyl and one methylmalonyl extender unit – again depending on whether the PKS belongs to the *trans*-AT or *cis*-AT class (Figure 9). In this scenario, the azumamide BGC would lack the amidase-encoding genes found in the gladiolin, etnangien and sorangicin BGCs. This would enable the azumamides to be produced as carboxamide-capped prodrugs, further diversifying the biological roles played by malonamyl starter units in polyketide biosynthesis.

### A unified view of amidated starter unit utilization by modular PKSs

Our work suggests that the malonamyl starter unit may be more widely employed by modular PKSs than hitherto appreciated (Figure 10). In addition, to serving as a key building block for the glutarimide pharmacophore in diverse *trans*-AT PKS products and the unique spirolactam in the sanglifehrins (assembled by a *cis*-AT PKS),^26^ it appears to be used as a “protected” malonyl derivative in sorangicin biosynthesis. An analogous role in gladiolin and etnangien biosynthesis is played by the sucinamyl starter unit, which may also be employed in sorangiolide biosynthesis. Moreover, a malonamyl starter unit may be utilized to mask the carboxylate-capped zinc-binding pharmacophore in the azumamides, enabling them to function as pro-drugs that are hydrolytically activated in target cells. Further experiments will be needed to substantiate this hypothesis.

**Figure 10.**
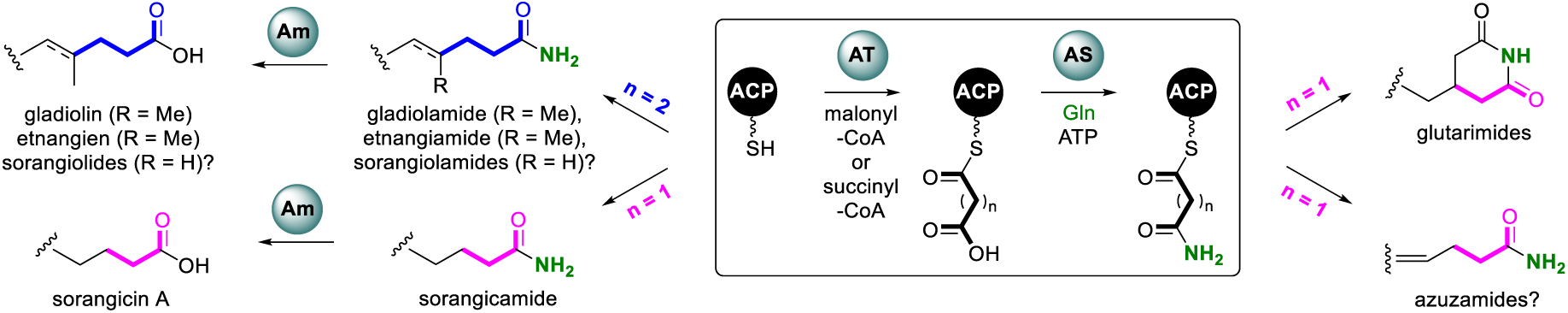
Unified mechanism for assembly of malonamyl and succinamyl starter units utilized by *trans*-AT PKSs. It remains to be substantiated whether this mechanism is employed in sorangiolide and azumamide biosynthesis.

In all *trans*-AT PKS-associated examples investigated to date, a unified biosynthetic mechanism is employed. This involves transfer of malonate or succinate from CoA to a dedicated ACP, followed by carboxyl group activation with ATP, and amidation using ammonia derived from glutamine hydrolysis (Figure 10).

### Prospects for biosynthetic engineering

The unified mechanism for assembly of malonamyl and succinamyl *trans*-AT PKS starter units suggests it may be possible to engineer the production of novel polyketide derivatives via a mix-and-match strategy. This notion is supported by the observation that different chain initiation mechanisms can coexist within a single microbial host, such as *B. gladioli* BCC1622, which produces gladiolin and gladiostatin from succinamyl and malonamyl starter units, respectively. Our observation of crosstalk between ATs and ACPs in the gladiolin and gladiostatin chain initiation modules provides preliminary evidence for the feasibility of this approach.

### Conclusions

Our work reveals an unanticipated mechanism for initiation of chain assembly by *trans*-AT PKSs, involving cryptic amidation of a succinyl starter unit in gladiolin **1** biosynthesis. Dissection of this mechanism using a combination of genetic, biochemical, and isotope labelling techniques showed it to be similar to that used by *trans*-AT PKSs for the incorporation of a malonamyl starter unit into structurally unrelated glutarimide antibiotics. Comparative bioinformatics and biosynthetic gene cluster reannotation led us to hypothesize that analogous mechanisms are employed to incorporate cryptic succinamyl and malonamyl starter units into etnangien and sorangicin A, respectively. Sorangicin A is structurally unrelated to either the glutarimide family of antibiotics or gladiolin / etnangien, suggesting succinamyl and malonamyl starter units may be incorporated into diverse polyketide structural classes. Retro-biosynthetic analyses suggest the sorangiolides may be a further polyketide structural class derived from a cryptic succinamyl starter unit and that the biosynthesis of the putative PKS-derived pharmacophore of azumamides family of tetrapeptidyl HDAC inhibitors may involve a malonamyl starter unit. Overall, our findings broaden our knowledge and deepen our understanding of chain initiation mechanisms in *trans*-AT PKSs and provide a foundation for engineered biosynthesis of novel polyketide derivatives containing rationally modified starter units.

## AUTHOR INFORMATION

### Author Contributions

‡Y.Z. and M.C. contributed equally to this work.

### Notes

The authors declare the following competing financial interest(s): G.L.C. is a co-founder, consultant, and shareholder of ErebaGen Ltd.

## ACKNOWLEDGMENTS

This research was supported by a PhD studentship to M.C. from the BBSRC Midlands Integrative Bioscience Doctoral Training Partnership (Grant Ref BB/J014532/1), a Chancellor’s International Scholarship to Y.Z. from the University of Warwick, and BBSRC research grants to G.L.C. (Grant Refs BB/M017982/1 and BB/W003171/1). We are grateful to Drs Lijiang Song and Ivan Prokes for assistance with obtaining MS and NMR data, respectively.

## TOC graphic

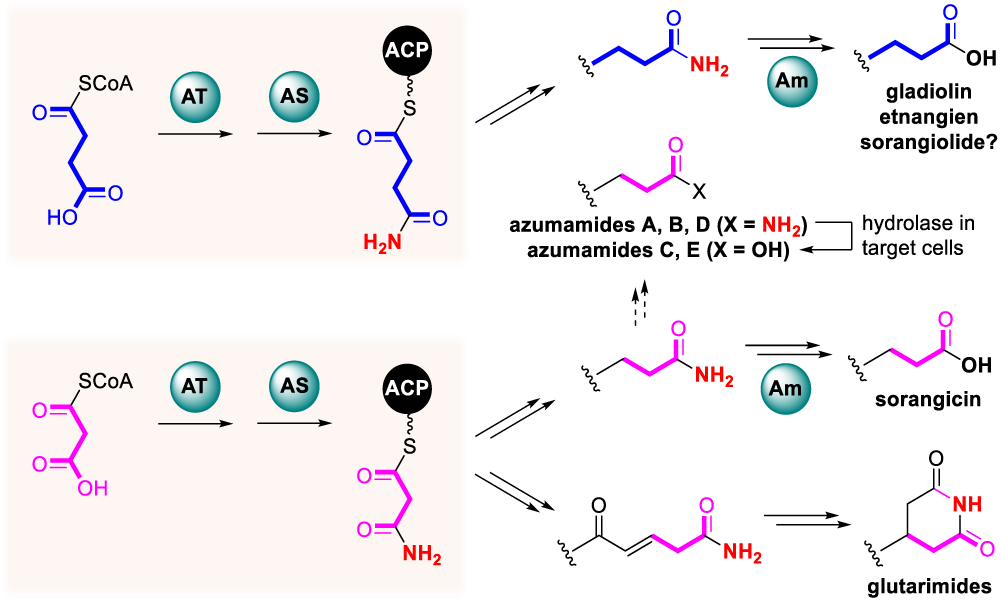

