## Supporting Information for "Cryptic starter amidation in antibiotic biosynthesis by *trans*-acyltransferase polyketide synthases"

### 1. Methods

#### 1.1 Gene deletion and complementation

In-frame deletions were created in *B. gladioli* BCC1622 using the suicide plasmid pGPI-*Sce*I and the I-*Sce*I nuclease expression plasmid pDAI-*Sce*I.[1] The pGPI-*Sce*I plasmid was constructed with 800-1000 bp homologous flanking sequences of each gene. The donor strain *E. coli* SY327, carrying pGPI-*Sce*I plasmid, was cultured along with overnight cultures of helper strain *E. coli* HB101 containing pRK2013 and *B. gladioli* BCC1622 for triparental mating. The cell pallets were harvested by centrifugation, suspended in 5 mL LB to remove the residue antibiotics, and then 100 μL of each resuspension in LB was mixed and spread onto a nitrocellulose disk (Millipore) placed on an antibiotic-free LB agar plate at 30°C overnight. The cell mixture was removed using 1 mL of sterile 0.9% NaCl and spread on LB agar with appropriate antibiotic at 30°C for 3 days. Exconjugants were selected for using 150 μg/mL trimethoprim and 600 U/mL polymyxin B and subjected to PCR screening to identify single crossover mutants. The procedure was repeated for the second homologous combination using the donor strain *E. coli* SY327 containing pDAI-*SceI*, the helper strain *E. coli* HB101 containing pRK2013, and the positive colony from the first homologous combination. Exconjugants were selected for using 200 μg/mL tetracycline and 600 U/mL polymyxin B and subjected to PCR screening to identify double crossover mutants. The removal of pDAI-*SceI* was achieved by growth on 15% sucrose to obtain the final mutants.

In *trans* genetic complementation was performed using the pMLBAD plasmid carrying the target gene.[2] The constructs were introduced into *E. coli* SY327 by electroporation and then transferred into the *B. gladioli BCC1622* mutants using triparental mating. Expression of each complemented gene was induced by 0.25% arabinose. As a control experiment, the empty pMLBAD vector was introduced to each deletion mutant. The production of gladiolin was analyzed using UHPLC-ESI-Q-ToF-MS.

##### 1.2 Gene cloning, protein overproduction and purification

The genes were amplified using Q5® Hot Start High-Fidelity DNA polymerase (New England Biolabs) with *B. gladioli* BCC1622 culture as the template and subsequently cloned into pET28a or pHis8_G2K vector containing N-terminal histidine tag. The recombinant construct was transformed into *E. coli* BL21(DE3) competent cells for protein expression. A single transformant was incubated in 10 mL LB as the seed culture, followed by subculturing into 1 L LB (37 °C, 180 rpm) until reaching $OD_{600}$ value of 0.6-0.8. The culture was induced with 0.4 mM IPTG and then incubated at 15 °C for 16-20 h for protein overproduction. The cell pellets were lysed using a constant system cell disrupter and then centrifugated at 17,000 rpm for 30 min. The resulting supernatant was loaded on a 1 mL HiTrap™ Chelating HP Column (GE HealthCare), pre-charged with 5 mL 0.1 M $NiSO_4$ and equilibrated with 10 mL loading buffer (20 mM Tris-HCl, 100 mM NaCl, 20 mM imidazole). The histidine-tagged proteins bound to the column were then eluted using a serial concentration (50 mM, 100 mM, 200 mM, 300 mM) of imidazole elution buffer. Fractions were analyzed by SDS-PAGE and the factions containing target proteins were collected and concentrated using a Vivaspin centrifuge filter (GE Healthcare) with an appropriate molecular weight cut off (MWCO) based on the size of the recombinant proteins.

##### 1.3 *In vitro* biochemical assays

###### 1.3.1 GbnM activity assay

This assay was conducted using cell lysate of *E. coli* BL21(DE3) carrying pET28a-*gbnM* or pET28a empty vector. Each transformant was incubated on a small scale (in 10 mL LB) for protein overproduction. IPTG was also added to the pET28a control for consistency. The cell pellets were harvested by centrifugation and then resuspended in 400 μL storage buffer (20 mM Tris-HCl, 100 mM NaCl, pH 8.0) prior to being lysed by sonication. Each of the lysates (5 μL) was mixed with gladiolamide dissolved in DMSO (2 μL, 800 μM) in a total volume of 50 μL with storage buffer. The reactions were incubated at RT for 3 h before adding 100 μL of MeCN to quench the reactions.

The precipitated proteins were removed by centrifugation and the supernatant were filtered using 0.45 µm nylon micro-centrifugal filters (Thermo Scientific) before high-resolution LC-MS analysis.

##### 1.3.2 AT domain loading assay

To investigate the substrate tolerance of GbnB, purified GbnB (100 µM) was incubated with acetyl-, malonyl, succinyl, glutaryl-, succinamic-CoA (800 µM) in storage buffer with final volume of 50 µL at RT for 10 mins. The resulting mixture was desalted using an Ultra-0.5 mL centrifugal filter with a 10 kDa MWCO membrane (Millipore) before intact protein MS analysis.

##### 1.3.3 Transacylation assay

The *apo*-ACP domain was first converted to active form using Sfp,[3] the phosphopantetheinyl (Ppant) transferase from *Bacillus subtilis*. The purified *apo*-ACP (200 µM) was treated with co-enzyme A (CoA, 2mM), Sfp (20 µM), $MgCl_2$ (10 mM) in storage buffer with total volume of 50 µL (RT 4 h for GbnA; RT 1 h for GbnM1ACP(I)). The excess CoA was removed by buffer exchanging using Ultra-0.5 mL centrifugal filters with a 3 kDa MWCO membrane (Millipore).

The transacylation assay was then conducted by incubating *holo*-ACP (100 µM) with purified AT domain (20 µM) and substrate (800 µM) in storage buffer at RT for 10 mins. A control reaction was performed using boiled AT domains. All samples underwent protein desalination by buffer exchange using Ultra-0.5 mL centrifugal filters a 3 kDa MWCO membrane (Millipore) before intact protein MS analysis.

#### 1.4 Antimicrobial activity assays

##### 1.4.1 Broth microdilution method

Overnight cultures of the test strains were grown in 5 mL Mueller Hinton media at 30 °C. The antibiotics were dissolved in DMSO to a concentration of 5 mg/mL. In a

96-well microtiter plate 50 μL of serial 2-fold dilutions of the antibiotic (128 μg/mL - 0.0025 μg/mL) in Mueller Hinton media were mixed with 50 μL of microbial suspension that was made up, from the overnight cultures, to McFarland standard No. 0.5 and then diluted 100-fold, also in Mueller Hinton media. The assays were incubated for 16 h at 30 °C, and the MIC was defined as the lowest concentration that visibly inhibited bacterial growth. Each assay was performed in triplicate.

##### 1.4.2 Resazurin microtiter assay

MICs against *Mycobacterium smegmatis* were determined using the rezasurin microtiter assay.[4] A culture of *M. smegmatis* was grown in mycobacteria media for 2 days at 37 °C while being shaken at 220 rpm. In a 96-well microtiter plate 50 μL of a bacterial suspension made up to McFarland standard No. 1, in 7H-9 media and diluted 20-fold was mixed with 50 μL of serial 2-fold dilutions of the antibiotic (128 μg/mL - 0.0025 μg/mL) also in 7H-9 media (initially the antibiotic was dissolved in DMSO to a concentration of 5 mg/mL). The assays were incubated at 37 °C for 2.5 days. 15 μL of resazurin solution (Acros Organics, made to a 0.01 % w/v solution in sterile water) was then added to each well and incubated overnight, the MIC was defined as the lowest concentration to prevent a complete color change from blue to pink. Each assay was performed in triplicate.

#### 1.5 Isotope labeling experiments

##### 1.5.1 Feeding experiment with [1,4-$^{13}C_2$]succinic acid to *B. gladioli* BCC1622

The glycerol stock of *B. gladioli* BCC1622 was cultured in 5 mL LB medium at 30 °C overnight. Cell pellets were obtained by centrifuging at 4,000 rpm for 5 mins, followed by the resuspension using 5 mL sterile 0.9% NaCl. The suspension was dipped with a cotton swab and streaked on the 3 mL BSM agar plate. At 0 h, a suspension of [1,4-$^{13}C_2$] succinic acid in DMSO was dropped onto the BSM plate to give a final concentration of 30 mM. After 36 h incubation at 30 °C, an equal volume of ethyl acetate (EtOAc) was added to each plate to extract the metabolites for 1 h and the extract was filtered and evaporated to concentrate for high-resolution LC-MS analysis. A scaled-up feeding experiment was conducted by adding 800 mg of [1,4-$^{13}C_2$]succinic acid to 800 mL BSM agar to give a final concentration of 8.3 mM. After 3 days of incubation, the $^{13}$C-lablleing metabolites were extracted using EtOAc.

##### 1.5.2 Feeding experiment with [amide-$^{15}$N]L-glutamine to *B. gladioli* BCC1622 Δ*gbnM*

The procedure was similar to the above feeding experiment with [1,4-$^{13}C_2$]succinic acid to *B. gladioli* BCC1622 except using the *gbnM* mutant strain. The suspension of harvested cell pellets of *B. gladoli* BCC1622 Δ*gbnM* in sterile 0.9% NaCl was dipped with a cotton swab and streaked on a 3 mL BSM agar plate. At 0 h, a suspension of [amide-$^{15}$N]L-glutamine in sterile water was dropped onto the BSM plate to give a final concentration of 20 mM. After incubation for 14 h, an equal volume of EtOAc was added to each plate to extract the metabolites for 1 h and the extract was filtered through a 0.4 μm filter (Thermo Scientific) and concentrated in vacuo for high-resolution LC-MS analysis.

#### 1.6 Isolation and characterization of gladiolamide and [$^{13}C_2$]gladiolin

Gladiolamide was obtained by culturing *B. gladioli BCC1622* Δ*gbnM* mutant on BSM agar plates at 30 °C for 3 days. A seed culture was prepared in 5 mL LB medium, and the cell pellets were harvested by centrifugation at 4,000 rpm for 5 min, followed by resuspension in 5 mL sterile 0.9% NaCl. The suspension was dipped with a cotton swab and streaked on the BSM agar plate containing glycerol as the sole carbon source. The surface cells were removed using a spreader, and the resulting agar was cut into small cubes using a spatula. Ethyl acetate was added in an equal volume to the agar cubes and left for 2 hours to extract the metabolites. The resulting extract was then filtered and concentrate in vacuo for subsequent HPLC purification.

The extract was purified using an Agilent 1260 Series HPLC instrument equipped with a HP Agilent 1260 Diode Array detector and a Thermo ScientificTM BetaSilTM C18 column (150 × 21.2 mm, 5 µm). The crude was dissolved in 2 mL 50% MeCN and then separated with an MeCN/$H_2O$ gradient increasing from 35% to 65% during the elution process, and the fractions containing the pure compounds were identified using HRMS. The desired fractions for gladiolamide were dried using a Genevac EZ-2 for NMR analysis.

[$^{13}C_2$]Gladiolin was extracted following the same procedure as for gladiolamide, except that the MeCN/$H_2O$ gradient used for HPLC purification was adjusted to increase from 40% to 70%.

#### 1.7 Mass spectrometry and NMR spectroscopy

##### 1.7.1 Intact proteins analysis by UHPLC-ESI-Q-ToF-MS

Purified proteins and *in vitro* assays were analyzed on a Bruker MaXis II ESI-Q-ToF-MS instrument connected to a Dionex 3000 RS UHPLC fitted with an ACE C4-300 RP column (100 x 2.1 mm, 5 µm, 30 °C). The column was eluted with a linear gradient of 5–100% MeCN containing 0.1% formic acid over 30 min. The mass spectrometer was operated in positive ion mode with a scan range of 200–3000 m/z. Source conditions were: end plate offset at −500 V; capillary at −4500 V; nebulizer gas ($N_2$) at 1.8 bar; dry gas ($N_2$) at 9.0 L min$^{-1}$; dry temperature at 200 °C. Ion transfer conditions were: ion funnel RF at 400 Vpp; multiple RF at 200 Vpp; quadrupole low mass at 200 m/z; collision energy at 8.0 eV; collision RF at 2000 Vpp; transfer time at 110.0 µs; pre-pulse storage time at 10.0 µs.

##### 1.7.2 Metabolite analysis by high-resolution MS

UHPLC-ESI-Q-ToF-MS analysis was performed using a Dionex UltiMate 3000 UHPLC connected to a Zorbax Eclipse Plus C-18 column (100 × 2.1 mm, 1.8 µm) coupled to a Bruker Compact mass spectrometer. The mobile phase was water and acetonitrile (MeCN), each supplemented with 0.1% formic acid with flow rate 0.2

mLmin$^{-1}$. The gradient profile was as follows: 0–5 mins 5% MeCN; 5–17 mins 5–100% MeCN; 17–22 mins 100% MeCN; 22–25 mins 100–5% MeCN; 25-34 mins 5% MeCN. The mass spectrometer was operated in positive-ion mode with a scan range of 50–3,000 m/z. Source conditions were: end-plate offset at −500 V, capillary at −4,500 V, nebulizer gas ($N_2$) at 1.6 bar, dry gas ($N_2$) at 81 min$^{-1}$ and dry temperature at 180 °C. Ion transfer conditions were: ion funnel radio frequency (RF) at 200 Vpp, multiple RF at 200 Vpp, quadrupole low mass at 55 m/z, collision energy at 5.0 eV, collision RF at 600 Vpp, ion cooler RF at 50–350 Vpp, transfer time at 121 μs and pre-pulse storage time at 1 μs. Calibration was performed with 1 mM sodium formate through a loop injection of 15 μL at the start of each run.

##### 1.7.3 Structure elucidation by NMR analysis

1 and 2D NMR spectra ($^1$H, $^{13}$C, COSY, HSQC, and HMBC) for structure elucidation were recorded using a Bruker Avance III 600 MHz instrument. The instrument operated at 600 MHz for $^1$H NMR and 150 MHz for $^{13}$C NMR with all spectra being recorded at 298 K. Purified gladiolamide (14 mg), *iso*-gladiolamide (5 mg), gladiolin(15 mg) and [$^{13}C_2$]gladiolin (15 mg) was dissolved in DMSO-$d_6$ for NMR analysis. Chemical shifts (δ) are given in ppm and coupling constants (J) are given in hertz (Hz).

#### 2. Figures

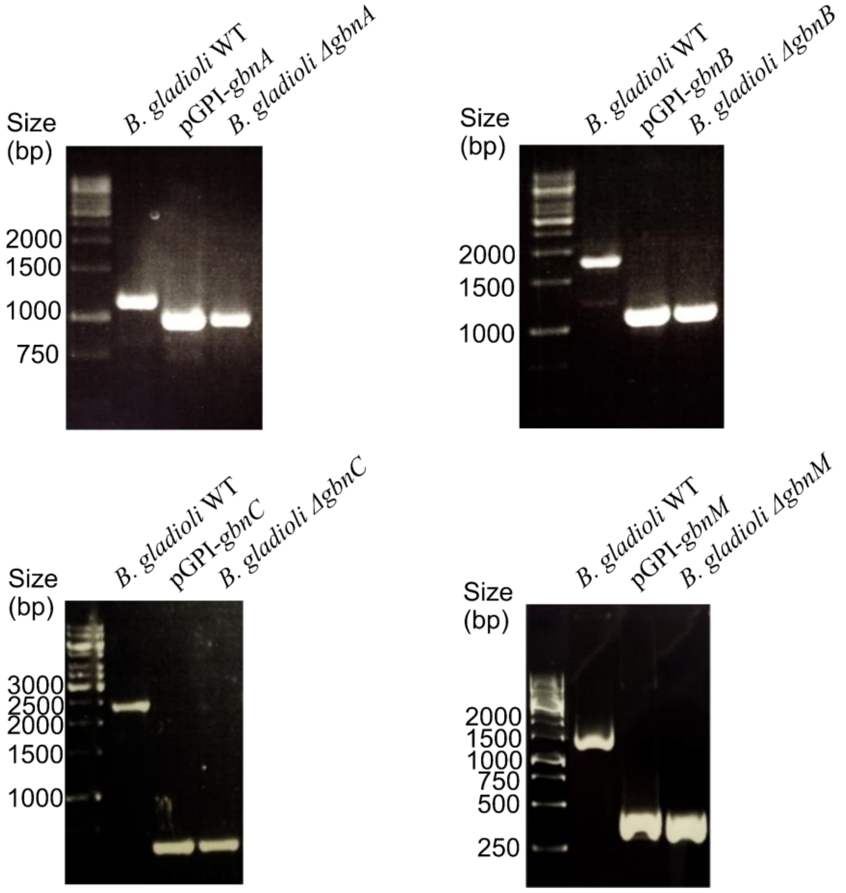


**Figure S1. PCR analysis confirming the construction of *gbnA, gbnB,* gbnC and *gbnM* mutants in *B. gladioli* BCC1622.** 1% agarose gel electrophoresis of PCR reactions confirm the creation of in-frame deletion in *gbnA*, *gbnB*, *gbnC* and *gbnM* with pGPI plasmid carrying the homologous flanking fragments as a positive control.

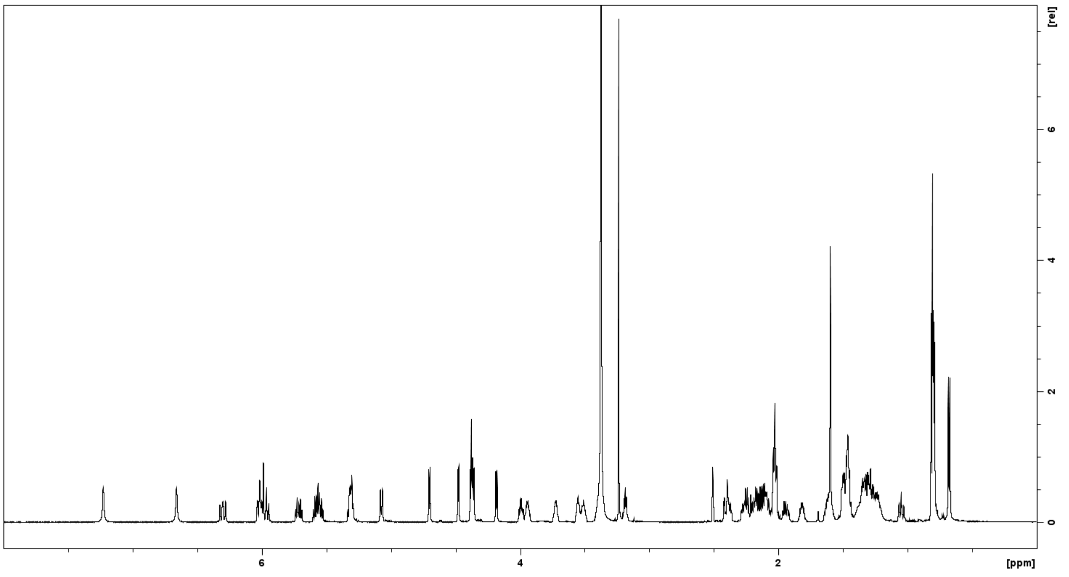


**Figure S2. $^{1}$H NMR spectrum of gladiolamide 2 in DMSO-$d_6$.**

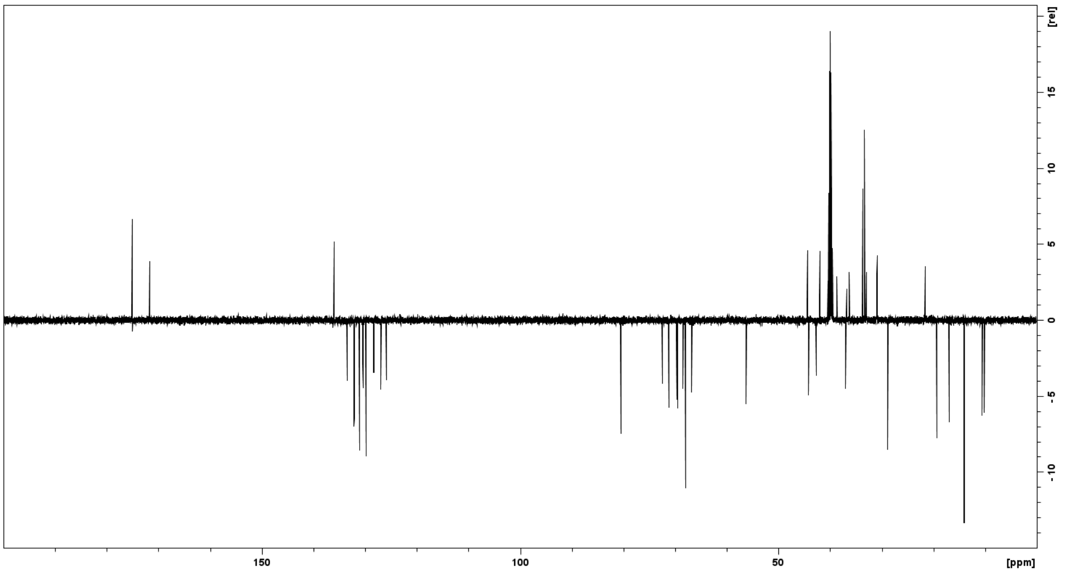


**Figure S3. $^{13}$C NMR spectrum of gladiolamide 2 in DMSO-$d_6$.**

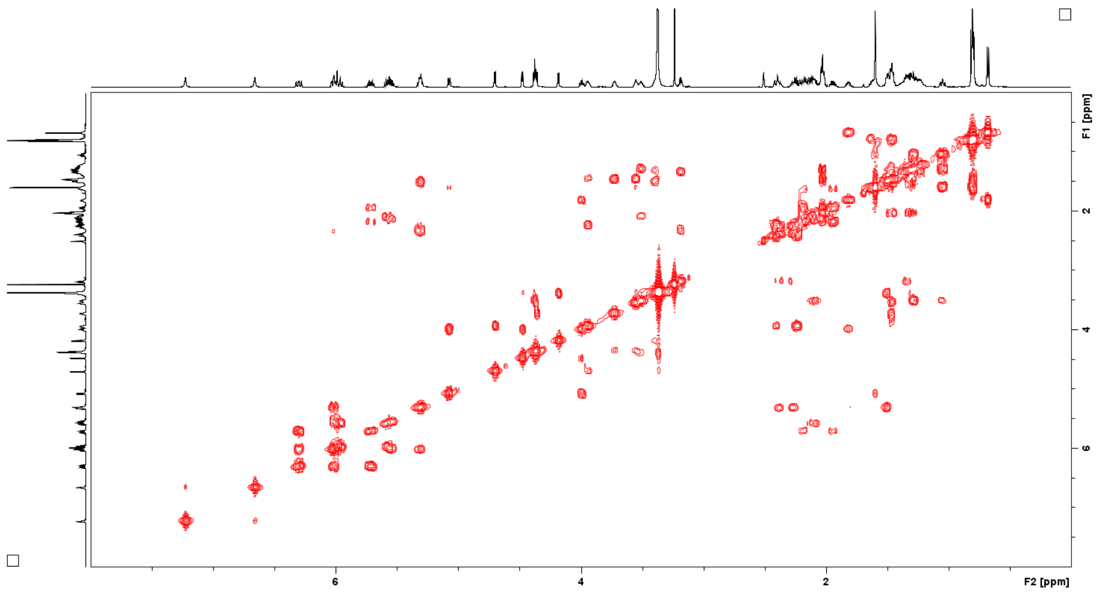


**Figure S4. COSY NMR spectrum of gladiolamide 2 in DMSO-d$_6$.**

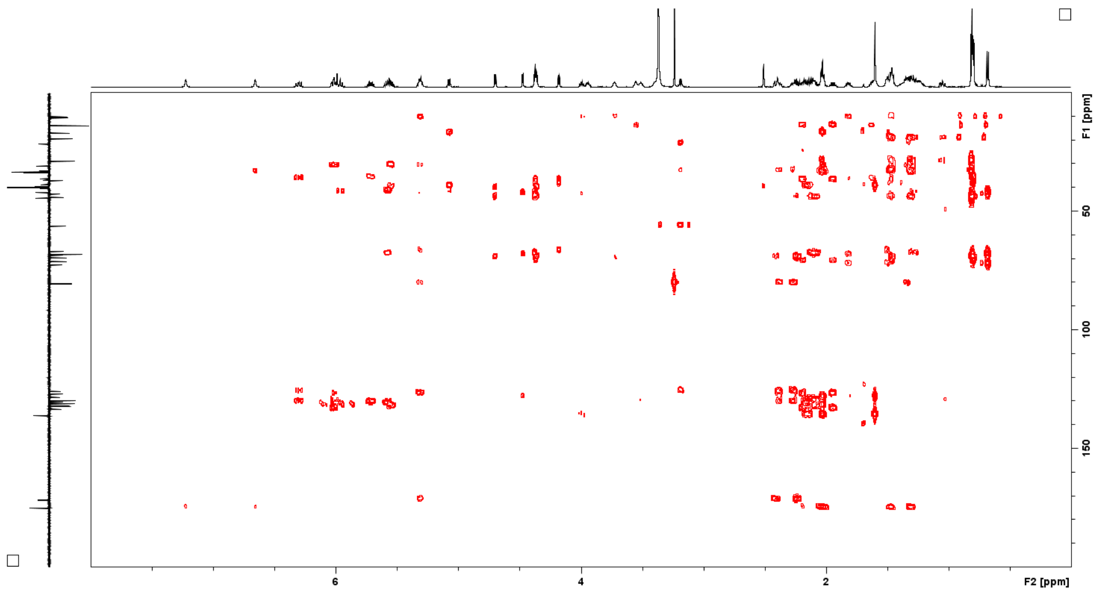


**Figure S5. HMBC NMR spectrum of gladiolamide 2 in DMSO-d$_6$.**

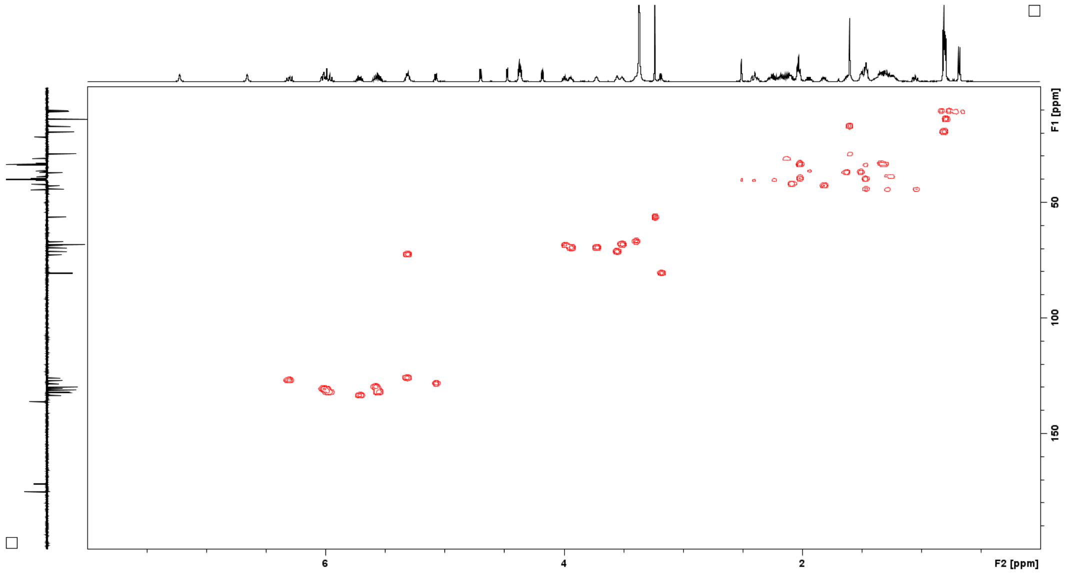


**Figure S6. HSQC NMR spectrum of gladiolamide 2 in DMSO-d6.**

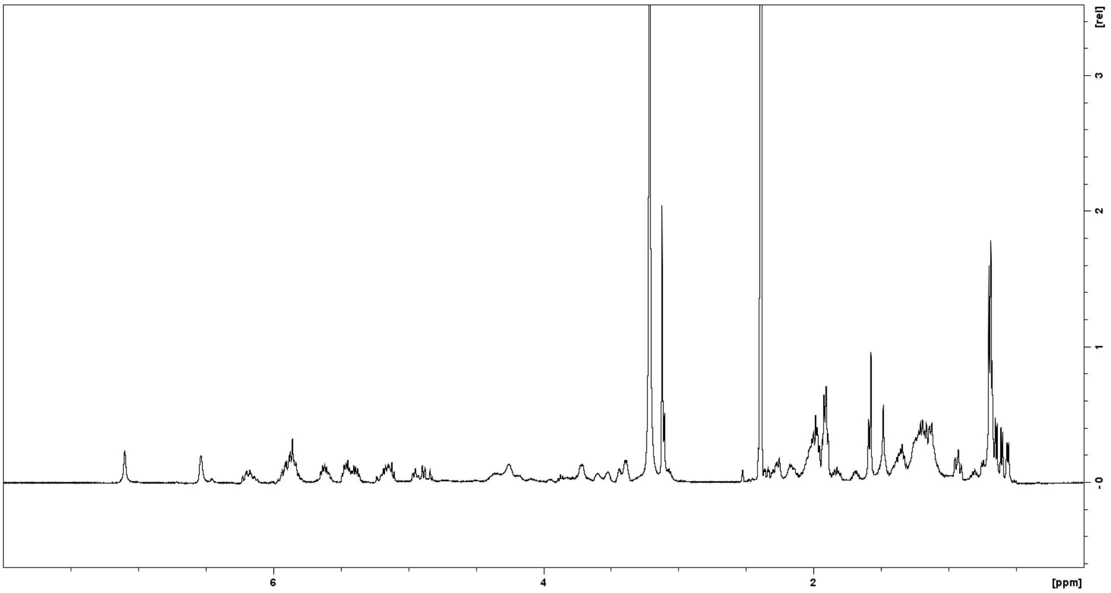


**Figure S7. [1]H NMR spectrum of *iso*-gladiolamide in DMSO-d6.**

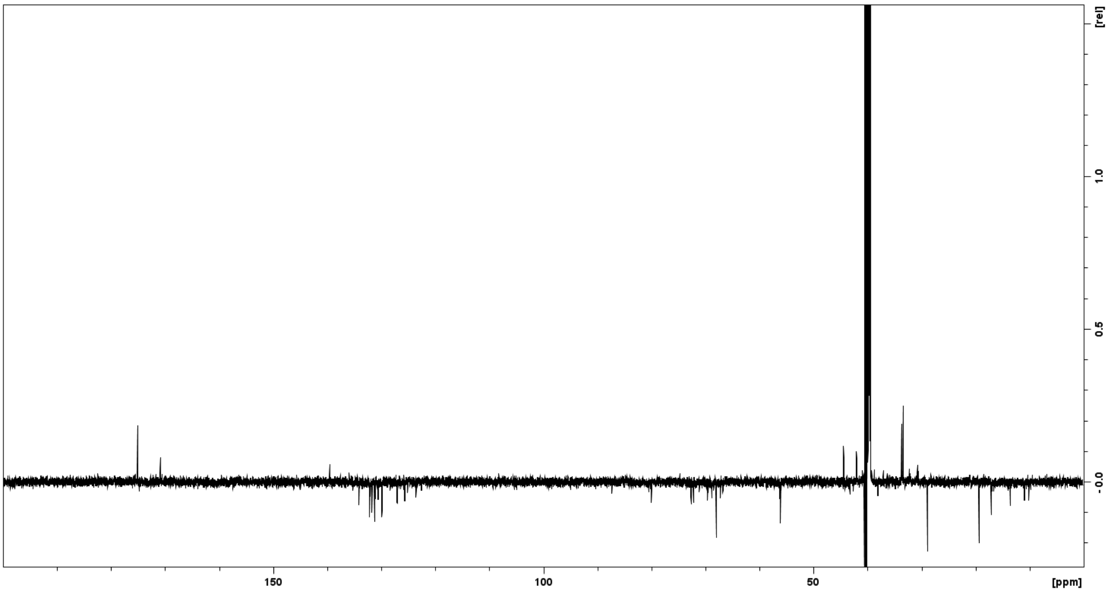


**Figure S8.** $^{13}$**C NMR spectrum of *iso*-gladiolamide in DMSO-d$_6$.**

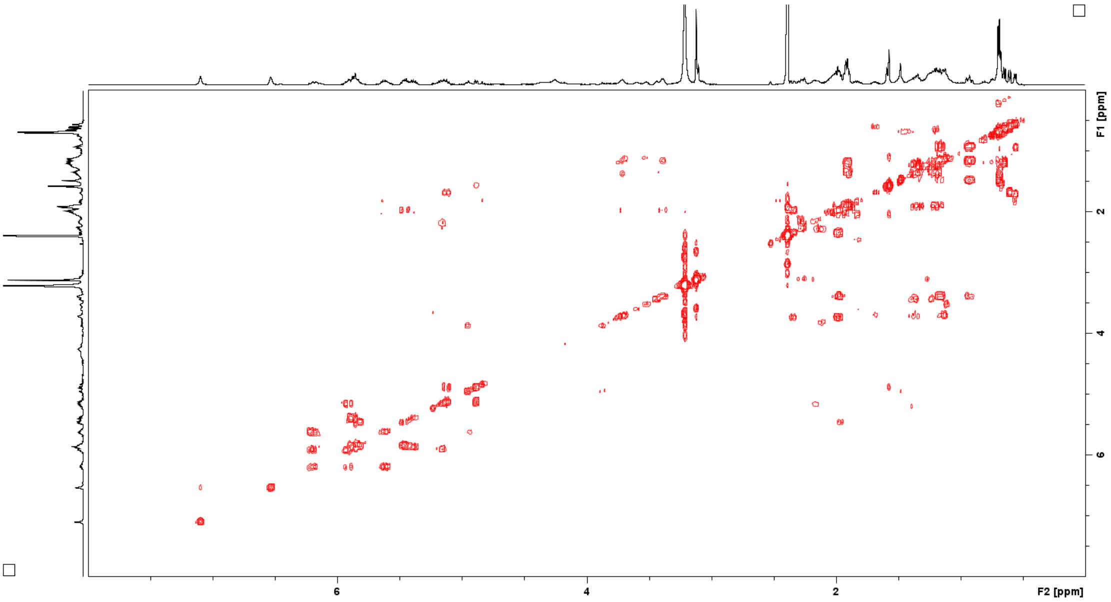


**Figure S9. COSY NMR spectrum of *iso*-gladiolamide in DMSO-d$_6$.**

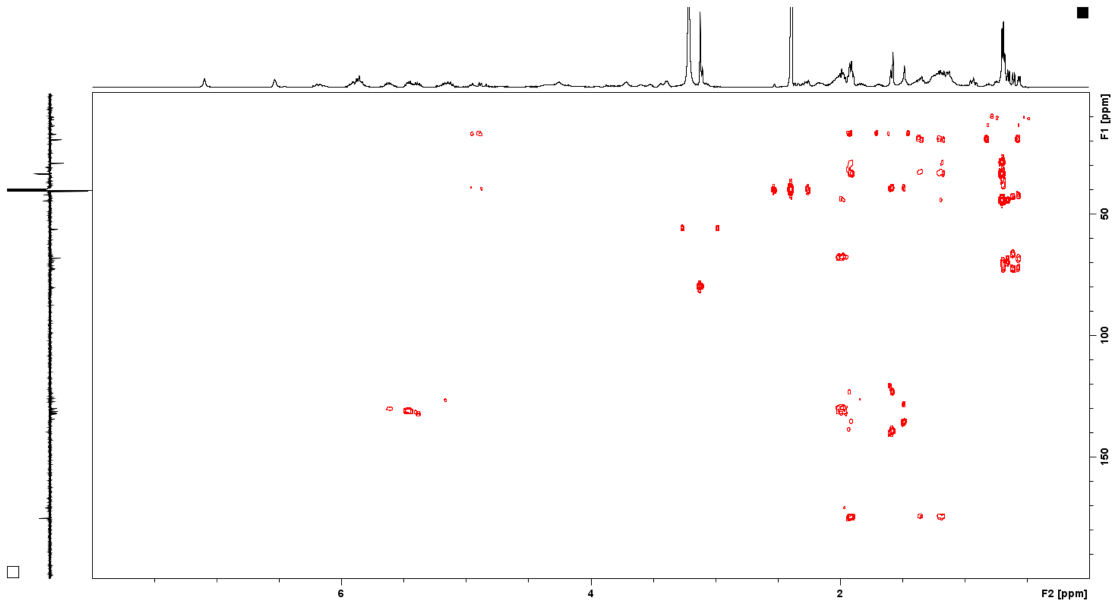


**Figure S10. HMBC NMR spectrum of *iso*-gladiolamide in DMSO-d$_6$.**

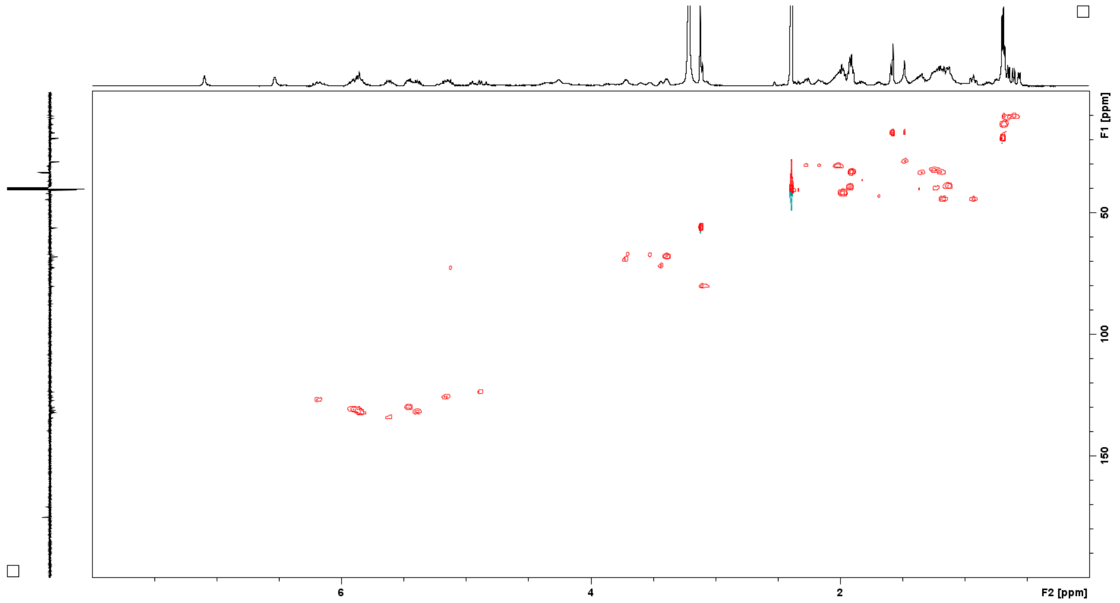


**Figure S11. HSQC NMR spectrum of *iso*-gladiolamide in DMSO-d$_6$.**

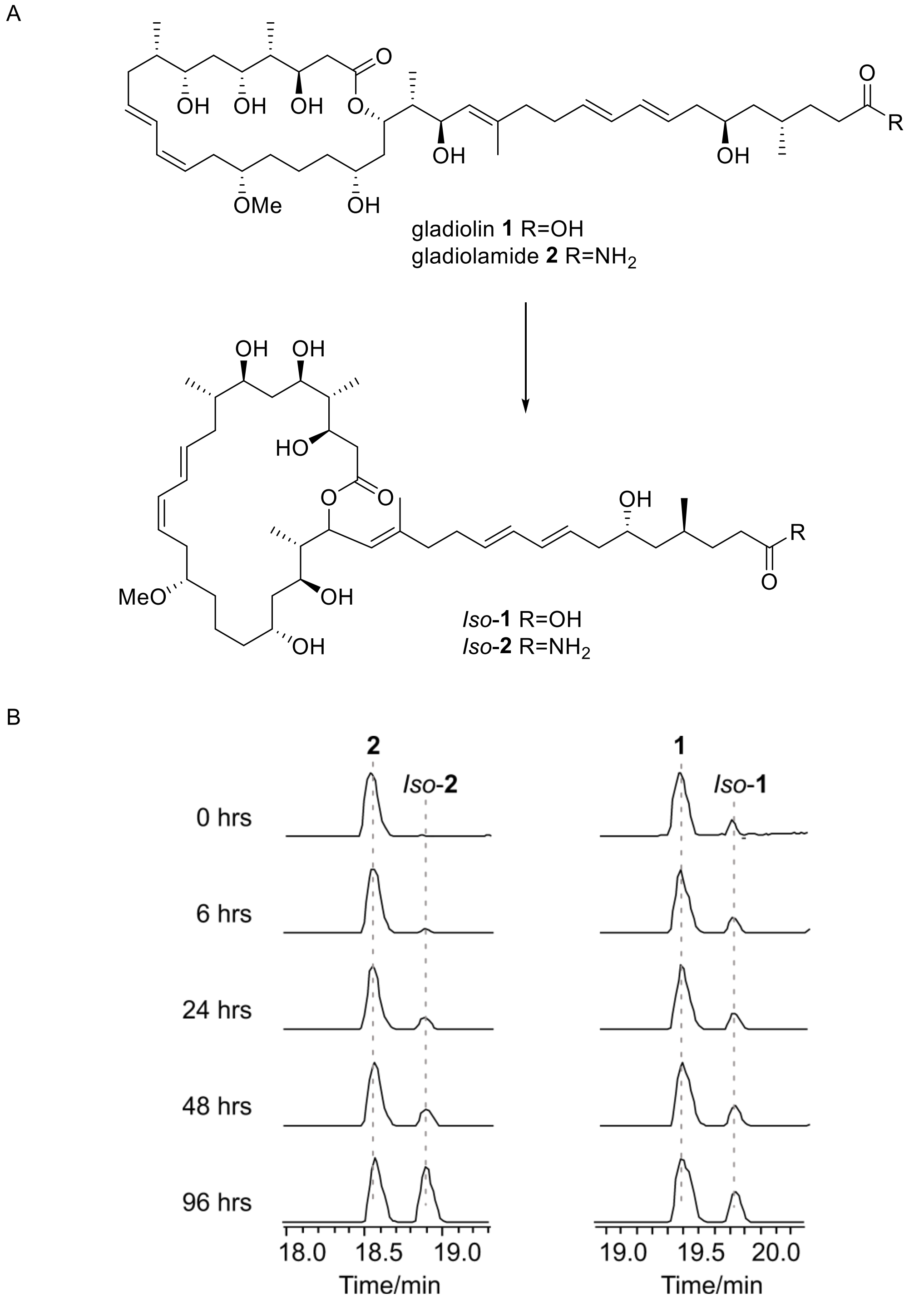


**Figure S12. The comparison for the conversion of gladiolin and gladiolamide to their isomers in MeOH.** (A) The proposed mechanism for the conversion of gladiolin and gladiolamide to their isomers with 24-membered lactone. (B) Base peak chromatograms from the time-course analysis showing the stability of gladiolin and gladiolamide in methanol.

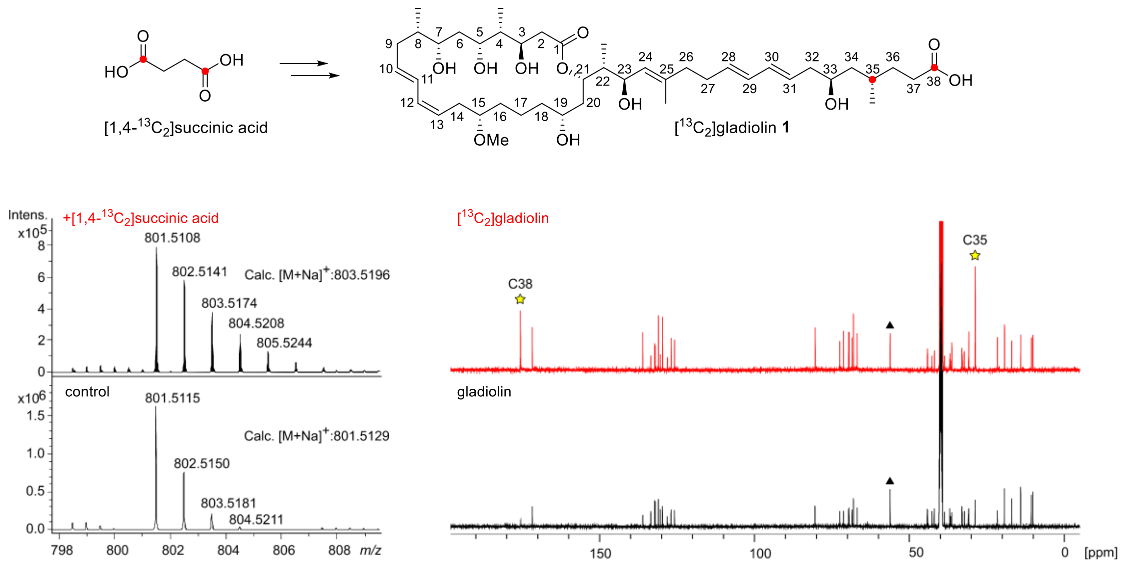


**Figure S13. Mass and NMR spectroscopic comparison of unlabeled gladiolin with [$^{13}C_2$]gladiolin derived from feeding of [1,4-$^{13}C_2$]succinic acid to *B. gladioli* BCC1622.** The increased peak with *m/z* = 803.5196 indicates the incorporation of [1,4-$^{13}C_2$]succinic acid in gladiolin biosynthesis. The signal due to the O-methyl group (labeled with a black triangle) was used to normalize the intensity of the NMR spectra. The signals due to C-35 and C-38 have considerably higher intensity in the spectrum of the labelled material (indicated by yellow stars).

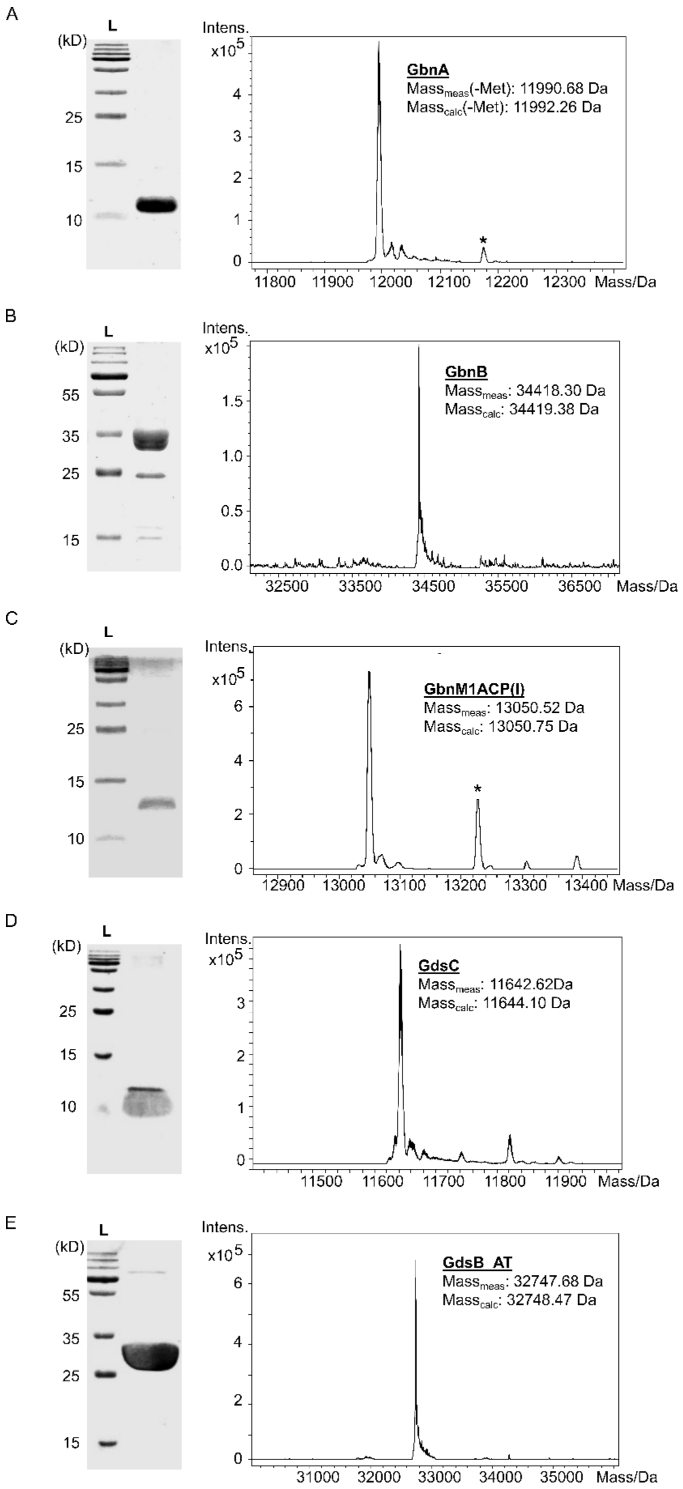


**Figure S14. SDS-PAGE analysis and deconvoluted ESI-MS spectra of purified N-His$_6$-GbnA, N-His$_8$-GbnB, N-His$_6$-GbnM1ACP(I), N-His$_8$-GdsC and N-His$_8$-GdsB_AT.** Peak labelled as '*' represents spontaneous gluconoylation of His$_6$-tag on fusion proteins, with an additional 178 Da observed.[5] The N-terminal Met residue of some proteins was removed by an internal aminopeptidase during overproduction in *E. coli* BL21 (DE3),[6] indicated as '(-Met)' in the calculated and measured masses.

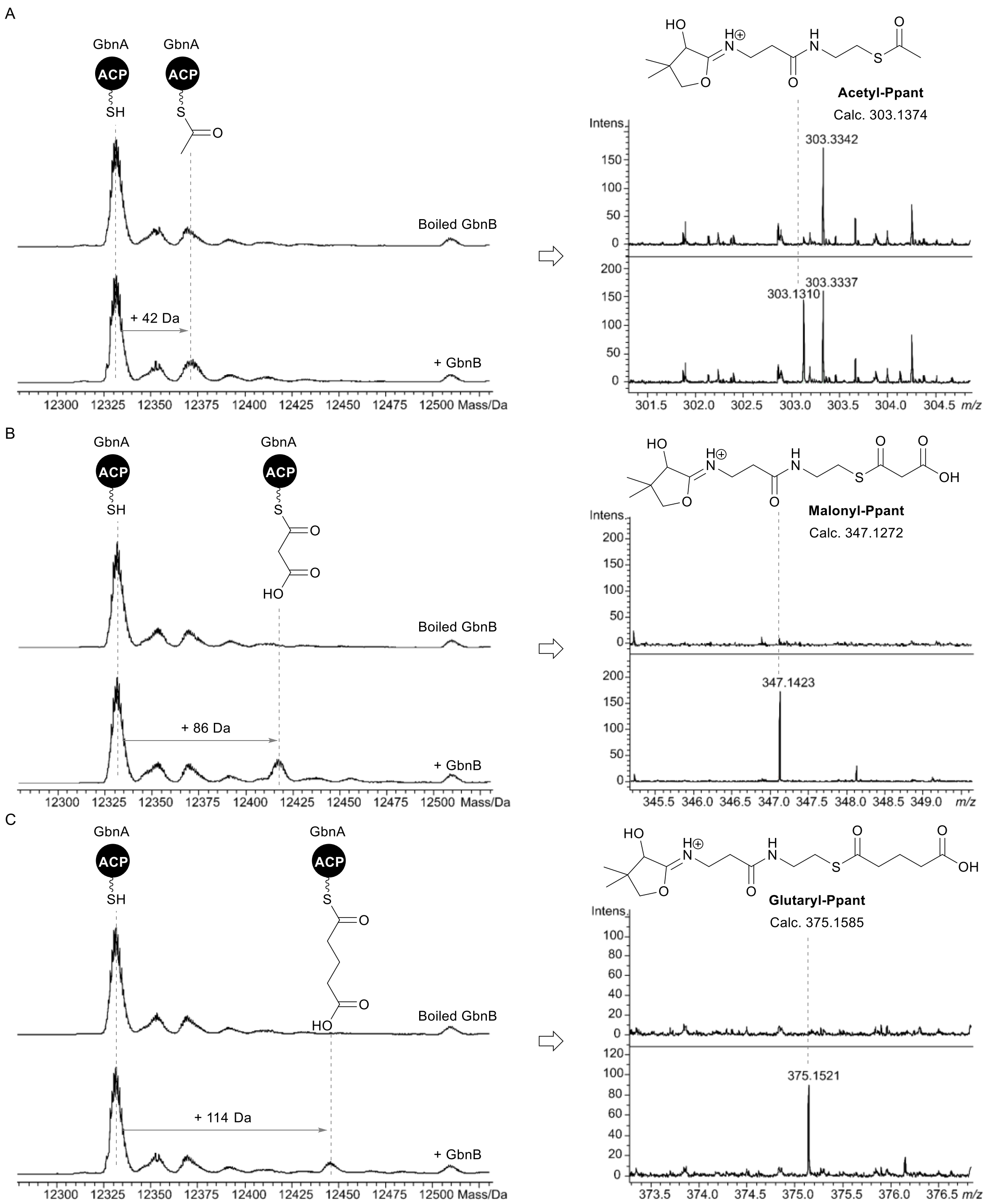


**Figure S15.** ***In vitro*** **transacylation assays showed the substrate tolerance of GbnB towards acetyl-CoA (A), malonyl-CoA (B) and glutaryl-CoA (C).** Deconvoluted mass spectra and PPant ejection analysis confirmed the transfer of acetyl-, malonyl- and glutaryl-CoA onto *holo*-GbnA by GbnB, evidenced by mass shift of 42 Da, 86 Da and 114 Da, respectively, in the mass spectra of the intact protein, and by the detection of ions with *m/z* = 303.1310, 347.1423 and 375.1521, corresponding to acetyl-, malonyl- and glutaryl-pantetheine.

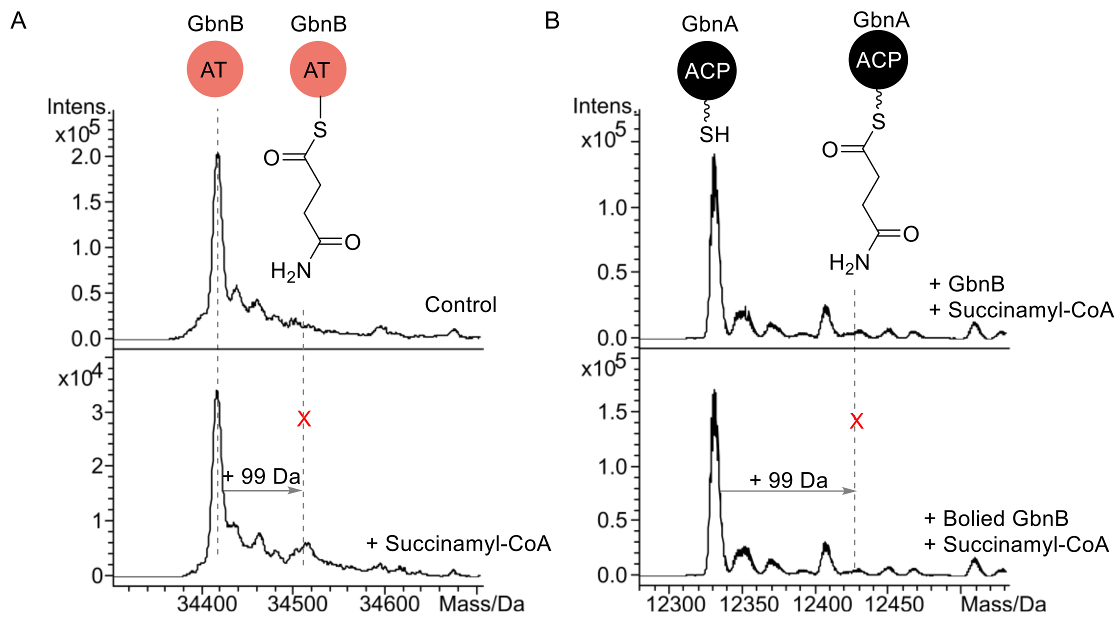


**Figure S16. *In vitro* assays showed that succinamyl-CoA is not the starter unit for gladiolin assembly.** The loading (A) and transacylation assays (B) involving GbnB, succinamyl-CoA, and *holo*-GbnA failed to generate either succinamyl-GbnB or succinamyl-GbnA, consistent with GbnC-catalyzed amidation of the succinyl unit after transfer onto GbnA.

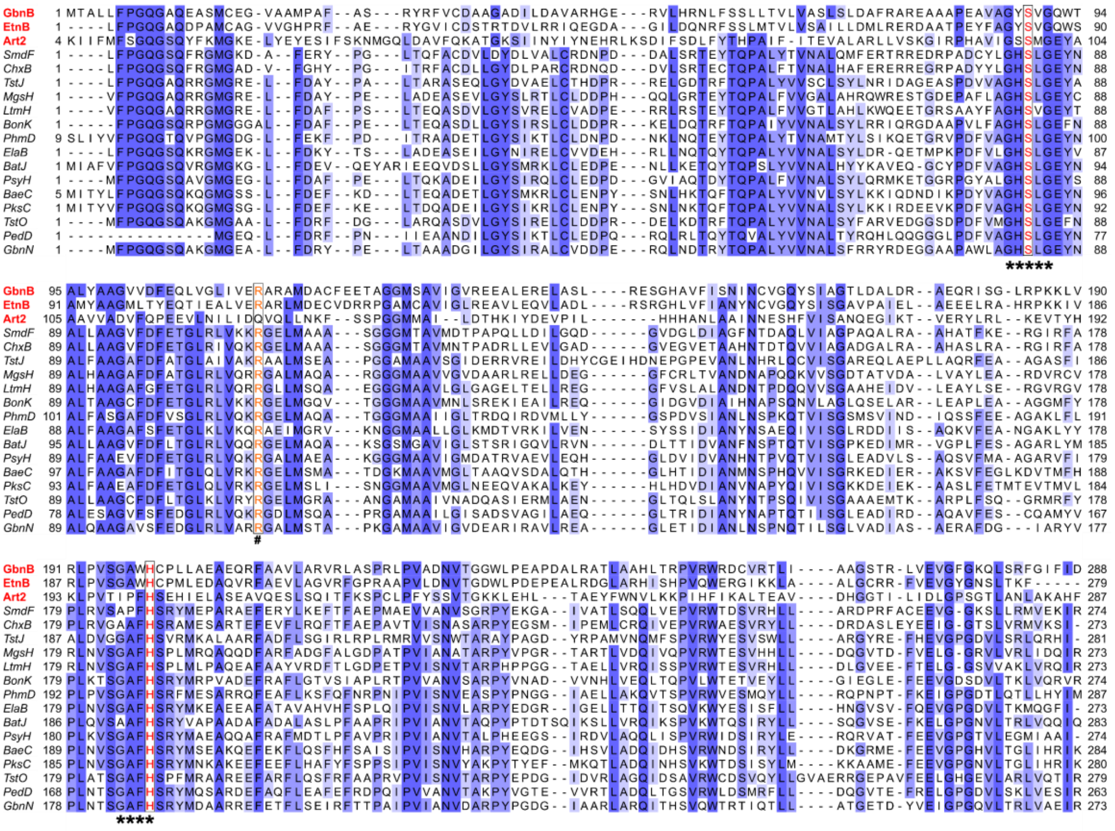

**Figure S17. Multiple sequence alignment of malonyl-specific and succinyl-specific *trans*-ATs.** Residues highlighted in red are part of the conserved Ser-His catalytic dyad. * indicates the conserved motifs in *trans*-AT domains: GHSLG and G(A)AFH motifs for malonyl-specific *trans*-ATs; GYSVG and GAWH motifs for succinyl-specific *trans*-AT domains. # indicates the conserved Arg in *trans*-ATs that is proposed to stabilize the carboxyl group of building blocks within the binding pocket, while it is substituted to Gln in acyl hydrolases,[7] as shown in Art2. Sequences from the following pathways were used: Etn, etnangien (BGC0000179); Art, aurantinins (BGC0001520); Smd, 9-methylstreptimidone (BGC0000171)； Chx, cycloheximide (BGC0000175); Tst, thailanstatin (BGC0001114); Mgs, migrastatin (BGC0000177); Ltm, lactimidomycin (BGC0000083); Bon, bongkrekic acid (BGC0000173); Phm, phormidolide (BGC0001350); Ela, elansolid (BGC0000178); Bat, batumin (BGC0001099); Psy, psymberin (BGC0001110); Bae/Pks, bacillaene (BGC0001089); Ped, pederin (BGC0001108).

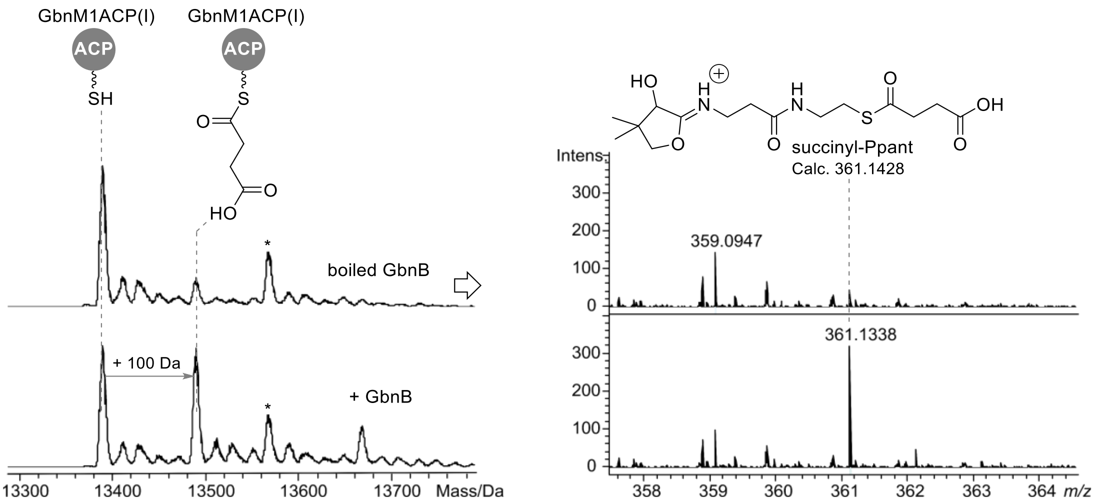


**Figure S18. *In vitro* examination of GbnB-catalyzed succinyl transfer onto GbnM1ACP(I).** Deconvoluted mass spectra (*left*) and Ppant ejection analysis (*right*) for the assays of GbnM1ACP(I) and GbnB, showing transfer of a succinyl unit onto *holo*-GbnM1ACP(I), evidenced by the mass shift of 100 Da in the mass spectrum and succinyl-PPant ion with *m/z* = 361.1338. Peak labelled as '*' represents spontaneous gluconoylation of $His_6$-tag on fusion proteins, with an additional 178 Da observed.[5]

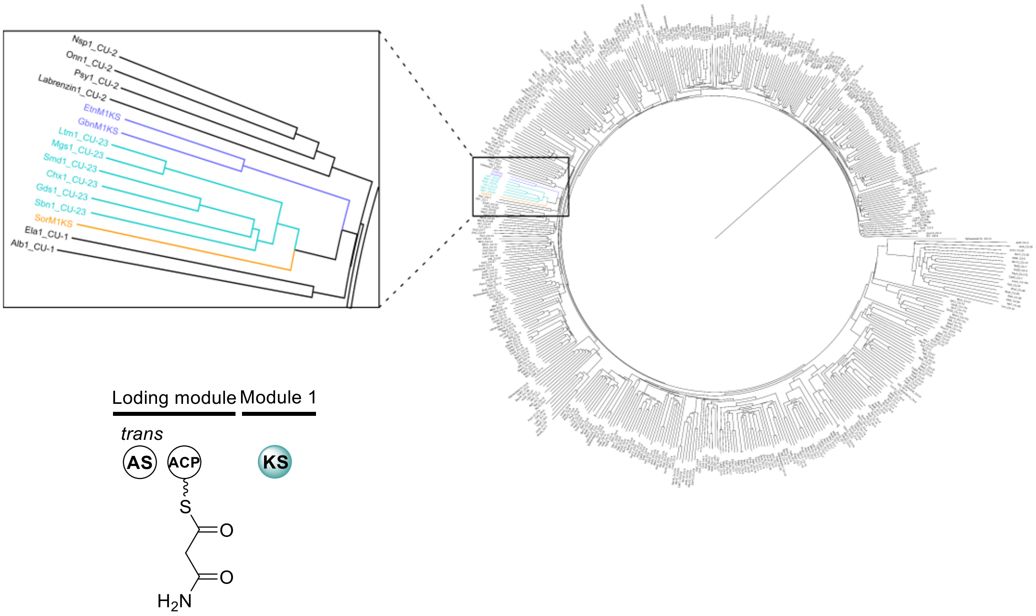


**Figure S19. Phylogenetic analysis of *trans*-AT PKS KS domains.** Module 1 KS domains from gladiolin, etnangien and sorangicin (GbnM1KS, EtnM1KS and SorM1KS, respectively) PKSs cluster tightly with those in several glutarimide PKSs (cyan), which are proposed to show substrate specificity for malonamyl units.

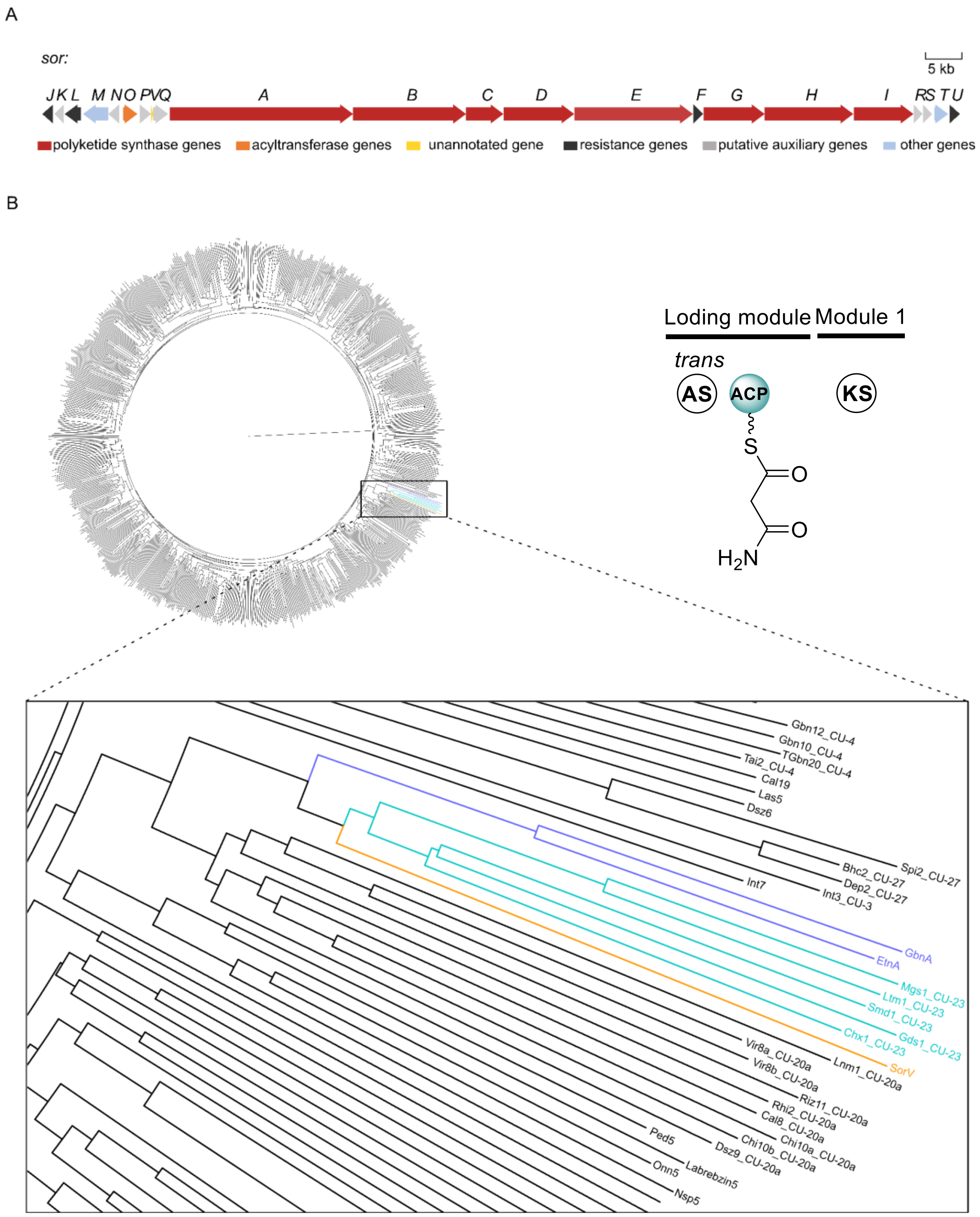


**Figure S20. Identification of a the previously overlooked *sorV* gene in the sorangicin BGC and phylogenetic analysis of SorV.** (A) Location of the previously overlooked gene, designated as *sorV*, encoding a standalone ACP. (B) Phylogenetic analysis of SorV shows that it clades with GbnA, EtnA and other homologs involved in glutarimide-containing polyketide (cyan) biosynthesis, which function as a platform for loading and amidation of malonyl-CoA.

### 3. Tables

**Table S1.** List of strains used in this study.

| Strain | Characteristic | Source/ Reference |
|---|---|---|
| ***Escherichia coli*** | | |
| Top10 | *mcrA,* Δ(*mrr-hsdRMS-mcrBC),* Phi80lacZ(*del)M15,* ΔlacX74*, deoR, recA1, araD139,* Δ(*ara-leu*) 7697, *galU, galK, rpsL(SmR), endA1, nupG,* for high-efficiency cloning and plasmid propagation. | Invitrogen |
| BL21(DE3) | *E. coli* str. B $F^-$ ompT gal dcm lon hsdSB($rB^-mB^-$) λ(DE3 [lacI lacUV5 T7p07 ind1 sam7 nin5]) [$malB^+$]$_{K-12}$($\lambda^S$), for expression and overproduction of recombinant proteins. | Invitrogen |
| SY327 | *araD, Δ(lac pro) argE(Am) recA56 rifR nalA λpir*, as a donor strain for conjugation between *E. coli* and *Burkholderia*. | Reference[8] |
| HB101 | *F- Lambda- araC14 leuB6(Am) DE(gpt-proA)62 lacY1 glnX44(AS) galK2(Oc) recA13 rpsL20(strR) xylA5 mtl-1 thiE1 hsdS20(rB*$^-$*, mB*$^-$*),* carries pRK2013, as a helper strain for conjugation between *E. coli* and *Burkholderia*. | Reference[9] |
| ***Burkholderia gladioli*** | | |
| BCC0238 | Wild type | Lab stock |
| BCC1622 | Wild type | Lab stock |
| BCC1622 Δ*gbnA* | *gbnA* mutant | This study |
| BCC1622 Δ*gbnA*::*gbnA* | *gbnA* complement | This study |
| BCC1622 Δ*gbnB* | *gbnB* mutant | This study |
| BCC1622 Δ*gbnB*::*gbnB* | *gbnB* complement | This study |
| BCC1622 Δ*gbnC* | *gbnC* mutant | This study |
| BCC1622 Δ*gbnC*::*gbnC* | *gbnC* complement | This study |
| BCC1622 Δ*gbnM* | *gbnM* mutant | This study |
| BCC1622 Δ*gbnM*::*gbnM* | *gbnM* complement | This study |

**Table S2.** List of vectors used in this study.

| Vector | Description | Source/Reference |
|---|---|---|
| pET28a | Bacterial vector for expression of N-terminally $His_6$-tagged proteins with a thrombin site. | Novagen |
| pHis8_G2K | The variant pET28a vector in which the second codon, Gly, is mutated to Lys to eliminate gluconoylation of purified proteins and the third and fourth codons, Ser, are mutated to His to form an N-termina $His_8$-tag. | Reference[10] |

| | | |
|---|---|---|
| pGPI-*SceI* | Suicide cloning vector designed to introduce a targeted I-SceI restriction site in the genome of *Burkholderia*. | Reference[1] |
| pDAI-*SceI* | Yeast homing endonuclease I-SceI expression vector. | Reference[1] |
| pMLBAD | Broad host range vector for arabinose-inducible in *trans* expression of proteins. | Reference[2] |

**Table S3.** List of primers used in this study.

Primers used for the creation of recombinant plasmids:

| **Plasmids** | **Primers (5'-3')/restriction site (NdeI + HindIII)** |
|---|---|
| pET28a-*gbnA* | For: AAAGGGCATATGCAAGACAAAATTCAGCAATTC<br>Rev: AAAGGGAAGCTTGGAACAGGAGCGCGGTCA |
| pHis8_G2K-*gbnB* | For: ATATCATATGACCGCGCTCCTGTTCCCC<br>Rev: CCCAAGCTTTCAATCGTAAGTGGTGCT |
| pET28a-*gbnC** | For: AAAAAACATATGGTGACCGTGTGCGGAATC<br>Rev: AAAAATAAGCTTTCATACCGTCACGCGTCC |
| pET28a-*gbnM1ACP(I)* | For: AAAGGGCATATGTTGCCGAAGGCCGCGGTC<br>Rev: TATGGGAAGCTTTCACGGTTCCGGAGCAGCCAG |
| pET28a-*gbnM** | For: AAAAAACATATGACTCCCGAACTCGACCTC<br>Rev: AAAAAAAAGCTTGCCGGCGTGTCAGTTGCC |
| pHis8_G2K-*gdsB*_AT | For: GGAATTCCATATGTACGCATTCGTTTTTCCTGG<br>Rev: CCCAAGCTTCTAGGCCTCGCTGCGGATCTTCGT |
| pHis8_G2K-*gdsC* | For: GGAATTCCATATGGAAAACCAGATTCGCGAG<br>Rev: CCCAAGCTTCTAGGCGAGCGCGACCTTGT |

* indicates that the plasmid expressed an insoluble protein.

Primers used for gene in-frame deletion and complementation:

| **Plasmids** | **Primers (5'-3')/restriction site** | |
|---|---|---|
| pGPI-*gbnA* | 5'-For: AAAGGGTCTAGATGCCAAGCGATGATGCGGTAA | XbaI |
| | 5'-Rev: AAAGGGAAGCTTGCGGTCCGACAGGAATTGCTG | HindIII |
| | 3'-For: AAGAGGAAGCTTCTGTCCGGAATCGTGTCGTTC | HindIII |
| | 3'-Rev: AGGAAGGGTACCTCAACTGCTTGCCGAAGCCGA | KpnI |
| | Screening-For: TCCAGATCGGATATTTCGGAA | |
| | Screening-Rev: TGATATTGCTGATGAACACCG | |
| pGPI-*gbnB* | 5'-For: AAAGGGTCTAGATTTTGATCGATAAGCTCGCGT | XbaI |
| | 5'-Rev: AAAGGGAAGCTTGAACCGATAACGGGATGCAAA | HindIII |
| | 3'-For: AAGAGGAAGCTTCGCTTGGTCGAGGTCGGCTTC | HindIII |
| | 3'-Rev: AAAGGGGGTACCCAGCCGCACGCTATCGGACAG | KpnI |
| | Screening-For: CGCGCCATGCAGTCATGATGG- | |
| | Screening-Rev: GAACAGCGGACGCTTGCCGAA- | |
| pGPI-*gbnC* | 5'-For: AAAGGGTCTAGAGATATTCTCGACGCCGTC | XbaI |
| | 5'-Rev: GGAGAACATATGCAGCATCCCCTTCAGGCA | NdeI |

| | | |
|---|---|---|
| | 3’-For: AAAGGGCATATGGAGAACCAGACCTTGTTG | NdeI |
| | 3’-Rev: AAAGGGGAATTCGGATCACGCCATACACCT | EcoRI |
| | Screening-For: TCGGCTTCGGCAAGCAGTTGA- | |
| | Screening-Rev: AGCAATCGGCATGGTCGATGA- | |
| pGPI-*gbnM* | 5’-For: ATATCTAGAGGCGGCAGGCTTCGTCTCGCG | XbaI |
| | 5’-Rev: ATAAAGCTTCGATTCGAGGTCGAGTTCGG | HindIII |
| | 3’-For: ATAAAGCTTATCGGCACGGTGCGAACG | HindIII |
| | 3’-Rev: ATAGGTACCGCGACCAGGCGTGTCAGG | KpnI |
| | Screening-For: GGCGTGGCCGGTGAAATCCT- | |
| | Screening-Rev: CGGCGGTCAGTTCCGGATA- | |
| pMLBAD-*gbnA* | For: AAAGGGCCATGGATGCAAGACAAAATTCA | NcoI |
| | Rev: AAAAAAAAGCTTTCATGCGCGCACGAAGCT | HindIII |
| pMLBAD-*gbnB* | For: AAAAAACCATGGATGACCGCGCTCCTGTT | NcoI |
| | Rev: AAAGGGAAGCTTTCAATCGTAAGTGGTGCT | HindIII |
| pMLBAD-*gbnC* | For: AAAGGGCCATGGGTGACCGTGTGCGGAATC | NcoI |
| | Rev: AAAAATAAGCTTTCATACCGTCACGCGTCC | HindIII |
| pMLBAD-*gbnM* | For: AAAAAACCATGGATGACTCCCGAACTCGACCTC | NcoI |
| | Rev: AAAAAAAAGCTTGCCGGCGTGTCAGTTGCC | HindIII |

**Table S4**. Comparison of [1]H NMR spectroscopic data observed for gladiolamide **2** isolated from *B. gladioli* BCC1622 Δ*gbnM* (DMSO-$d_6$, 600 Hz) and gladiolin published data. * indicates that the signal was found under the DMSO-$d_6$ peak, and ** indicates that the signal was found under the water peak. Highlighted in blue are the chemical shift assignments that differ most significantly. Multiplicity is denoted as follows: s = singlet, d = doublet, dd = doublet of doublets, dt = doublet of triplets, br = broad, m = multiplet and ol = overlapped.

| **Position** | **Gladiolamide** | | **Gladiolin** | |
|---|---|---|---|---|
| | $\delta_H$ (*J*, Hz) | $\delta_C$ | $\delta_H$ (*J*, Hz) | $\delta_C$ |
| 1 | - | 171.3 | - | 170.9 |
| 2 | 2.22, 2.40, 2 x1H, 2 x m | 39.6* | 2.22, 2.40, 2 x1H, 2 x m | 39.6 |
| 3 | 3.94, 1H, m | 69.3 | 3.95, 1H, m | 68.9 |
| 3-OH | 4.69, 1H, d, (5.6) | - | 4.05, ol | - |
| 4 | 1.46, 1H, m | 43.7 | 1.46, 1H, m | 43.8 |
| 4-Me | 0.79, 3H, d, (6.4) | 9.7 | 0.79, 3H, d, (6.5) | 9.6 |
| 5 | 3.72, 1H, m | 69.1 | 3.71, 1H, m | 68.8 |
| 5-OH | 4.35, 1H, ol | - | 4.02, ol | - |
| 6 | 1.46, 2H, m, ol | 39.2* | 1.46, 2H, m | 39.6 |
| 7 | 3.54, 1H, m | 70.7 | 3.54, 1H, m | 70.6 |

| | | | | |
|---|---|---|---|---|
| 7-OH | 4.36, 1H, ol | - | 4.02, 1H, br s | - |
| 8 | 1.62, 1H, m | 36.5 | 1.62, 1H, m | 37.0 |
| 8-Me | 0.79, 3H, d, (6.7) | 13.6 | 0.78, 3H, d, (6.0) | 13.3 |
| 9 | 1.94, 2.18, 2 x 1H, 2 x m | 35.9 | 1.92, 2.18, 2 x 1H, 2 x m | 35.8 |
| 10 | 5.71, 1H, dt, (6.6, 14.5) | 133.0 | 5.71, 1H, dt, (6.5, 15.0) | 133.0 |
| 11 | 6.29, 1H, dd, (10.96, 15.0) | 126.5 | 6.29, 1H, dd, (11.0, 15.0) | 126.1 |
| 12 | 6.01, 1H, ol | 130.0 | 6.02, 1H, (11.0, 11.0) | 129.7 |
| 13 | 5.31, 1H, ol | 125.4 | 5.31, 1H, dt, (7.0, 11.0) | 125.0 |
| 14 | 2.26, 2.36, 2 x 1H, 2 x m | 30.6 | 2.26, 2.38, 2 x 1H, 2 x m | 30.0 |
| 15 | 3.18, 1H, m | 80.1 | 3.18, 1H, dd, (5.5, 5.5) | 79.6 |
| 15-OMe | 3.23, 3H, s | 55.8 | 3.23, 3H, s | 55.5 |
| 16 | 1.31, 1.46, 2 x 1H, 2 x m | 33.2 | 1.32, 1.48, 2 x 1H, 2 x m | 32.2 |
| 17 | 1.21, 1.36, 2 x 1H, 2 x m | 21.2 | 1.23, 1.36, 2 x 1H, 2 x m | 21.1 |
| 18 | 1.24, 1.28, 2 x 1H, m, ol | 38.3 | 1.32, 1.30, 2 x 1H, 2 x m | 36.2 |
| 19 | 3.38** | 66.4 | 3.38, 1H, m | 66.1 |
| 19-OH | 4.17, 1H, d, (5.9) | - | 3.54, 1H, ol | - |
| 20 | 1.50, 2H, m | 36.4 | 1.49, 1.61, 2 x 1H, 2 x m | 36.3 |
| 21 | 5.30, 1H, m, ol | 72.0 | 5.30, 1H, m | 71.8 |
| 22 | 1.80, 1H, m | 42.2 | 1.80, 1H, m | 42.1 |
| 22-Me | 0.67, 3H, d, (7.03) | 10.2 | 0.69, 3H, d, (6.5) | 10.4 |
| 23 | 3.99, 1H, m | 68.1 | 3.98, 1H, m | 67.8 |
| 23-OH | 4.66, 1H, d, (4.03) | - | 4.46, 1H, br s | - |
| 24 | 5.06, 1H, d, (8.90) | 127.9 | 5.06, 1H, d, (8.50) | 127.6 |
| 25 | - | 135.5 | - | 134.7 |
| 25-Me | 1.59, 3H, s | 16.5 | 1.59, 3H, s | 16.2 |
| 26 | 2.02, 2H, ol | 39.1* | 2.02, 2H, m | 38.9 |
| 27 | 2.14, 2H, m | 30.5 | 2.12, 2H, m | 30.1 |
| 28 | 5.54, 1H, ol | 131.6 | 5.54, 1H, dt, (7.0, 15.0) | 130.2 |
| 29 | 5.98, 1H, m, ol | 130.6 | 5.98, 1H, dd, (11.0, 15.0) | 131.1 |
| 30 | 5.95, 1H, ol | 131.7 | 5.96, 1H, dd, (11.0, 15.0) | 131.9 |
| 31 | 5.57, 1H, ol | 129.4 | 5.56, 1H, dt, (7.0, 15.0) | 129.2 |
| 32 | 2.08, 2H, m | 41.6 | 2.08, 2H, m | 41.3 |
| 33 | 3.99, 1H, m | 67.6 | 3.49, 1H, m | 67.1 |
| 33-OH | 4.37, 1H, ol | - | 3.80, 1H, br s | - |
| 34 | 1.04, 1.28, 2 x 1H, 2 x m | 44.0 | 1.04, 1.28, 2 x 1H, 2 x m | 43.6 |
| 35 | 1.61, 1H, m, ol | 28.4 | 1.61, 1H, m | 28.1 |
| 35-Me | 0.80, 3H, d, (6.3) | 18.9 | 0.80, 3H, d, (6.5) | 17.9 |
| 36 | 2.02, 2H, m, ol | 32.5 | 1.32, 1.47, 2 x 1H, 2 x m | 32.3 |
| 37 | 1.29, 1.46, 2 x 1H, 2 x m | 32.9 | 2.14, 2.19, 2 x 1H, 2 x m | 29.9 |
| 38 | - | 174.7 | - | 174.9 |
| $NH_2$ | 6.65, 7.22, 2 x 1H, 2 x br s | - | - | - |

**Table S5.** $^{1}$H NMR data for *iso*-gladiolamide (DMSO-d$_6$, 600 Hz). * indicates that the signal was found under the DMSO-d$_6$ peak. Multiplicity is denoted as follows: s = singlet, d = doublet, dd = doublet of doublets, dt =doublet of triplets, br = broad, m = multiplet and ol = overlapped.

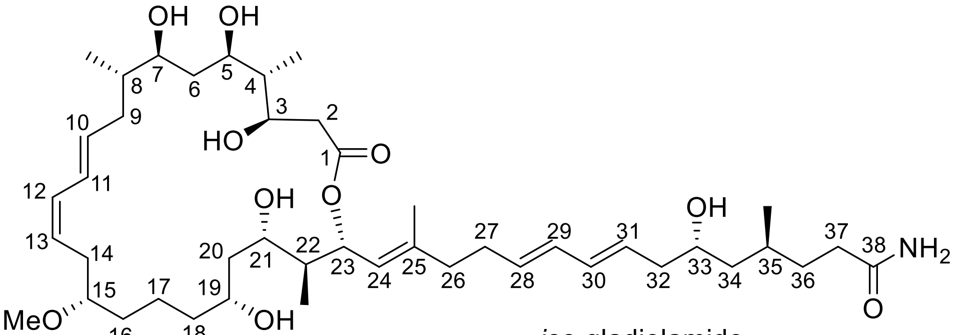


*iso*-gladiolamide

| Position | $\delta_H$ (*J*, Hz) | $\delta_C$ |
|---|---|---|
| **1** | - | 170.41 |
| **2** | 2.15, 2.45, 2 x 1H, 2 x m | 39.6* |
| **3** | 3.72, 1H, m | 68.35 |
| **4** | 1.45, 1H, m | 43.5 |
| **4-Me** | 0.76, 3H, d, (6.9) | 9.7 |
| **5** | 3.84, 1H, m | 69.1 |
| **6** | 1.35, 1.48, 2H, m, ol | 39.3* |
| **7** | 3.55, 1H, m | 71.7 |
| **8** | 1.51, 1H, m | 37.7 |
| **8-Me** | 0.79, 3H, d, (6.7) | 13.1 |
| **9** | 1.93, 2.16, 2 x 1H, 2 x m | 36.5 |
| **10** | 5.73, 1H, dt, (6.8, 14.3) | 133.6 |
| **11** | 6.31, 1H, dd, (11.0, 14.8) | 126.57 |
| **12** | 6.01, 1H, ol | 130.1 |
| **13** | 5.27, 1H, ol | 125.2 |
| **14** | 2.28, 2.38, 2 x 1H, 2 x m | 30.1 |
| **15** | 3.21, 1H, m | 79.6 |
| **15-OMe** | 3.23, 3H, s | 55.7 |
| **16** | 1.36, 1H, m | 31.8 |
| **17** | 1.22, 1.36, 2 x 1H, 2 x m | 20.6 |
| **18** | 1.24, 1.32, 2H, m, ol | 38.2 |
| **19** | 3.64 | 66.8 |
| **20** | 1.35, 1.47, 2 x 1H, 2 x m | 39.3 |
| **21** | 3.82, 1H, m | 66.3 |
| **22** | 1.80, 1H, m | 42.2 |

| | | |
|---|---|---|
| **22-Me** | 0.71 3H, d, (7.0) | 10.6 |
| **23** | 5.24, 1H, m | 72.2 |
| **24** | 5.00, 1H, d, (9.2) | 123.1 |
| **25** | - | 139.1 |
| **25-Me** | 1.69, 3H, s | 16.6 |
| **26** | 2.03, 2H, ol | 39.06* |
| **27** | 2.13, 2H, m | 30.3 |
| **28** | 5.51, 1H, m | 131.3 |
| **29** | 5.96, 1H, m, ol | 130.7 |
| **30** | 5.96, 1H, ol | 131.7 |
| **31** | 5.57, 1H, m | 129.4 |
| **32** | 2.09, 2H, m | 41.6 |
| **33** | 3.50, 1H, m | 67.5 |
| **34** | 1.04, 1.28, 2 x 1H, 2 x m | 43.9 |
| **35** | 1.60, 1H, m, ol | 28.4 |
| **35-Me** | 0.80, 3H, d, (6.5) | 18.9 |
| **36** | 2.02, 2H, m, ol | 32.5 |
| **37** | 1.30, 1.46, 2 x 1H, 2 x m | 32.9 |
| **38** | - | 174.5 |
| **$NH_2$** | 6.65, 7.21, 2 x 1H, 2 x br s | - |

**Table S6.** Comparison of the antimicrobial activity of *iso*-gladiolamide and gladiolamide (MIC = minimum inhibitory concentration).

| **Strains** | **MIC [μgmL$^{-1}$]** | |
|---|---|---|
| | ***Iso*-gladiolamide** | **Gladiolamide** |
| Gram-negative bacteria | | |
| *Klebsiella pneumonia* DSM26371 | >64 | >64 |
| *Acinetobacter baumannii* DSM25645 | 16 | 8 |
| *Pseudomonas aeruginosa* DSM29239 | >64 | 64 |
| *Enterobacter cloacae* DSM16690 | >64 | 64 |
| *Escherichia coli* SY327 | 32 | 16 |
| *Burkholderia gladioli* BCC1622 | >64 | >64 |
| Gram-positive bacteria | | |
| *Enterococcus faecium* DSM25390 | 2 | 2 |
| *Staphylococcus aureus* DSM21979 | 8 | 4 |
